# Selective engagement of prefrontal VIP neurons in reversal learning

**DOI:** 10.1101/2024.04.03.587891

**Authors:** Jee Hyun Yi, Young Ju Yoon, Huijeong Jeong, Seo Yeon Choe, Min Whan Jung

**Author notes:** These authors contributed equally to this work (Jee Hyun Yi, Young Ju Yoon).

## Abstract

To gain insights into neural mechanisms enabling behavioral adaptations to complex and multidimensional environmental dynamics, we examined roles of VIP neurons in mouse medial prefrontal cortex (mPFC) in probabilistic reversal learning. Behaviorally, manipulating VIP neuronal activity left probabilistic classical conditioning unaffected but severely impaired reversal learning. Physiologically, conditioned cue-associated VIP neuronal responses changed abruptly after encountering an unexpected reward. They also conveyed strong reward prediction error signals during behavioral reversal, but not before or after, unlike pyramidal neurons which consistently conveyed error signals throughout all phases. Furthermore, the signal’s persistence across trials correlated with reversal learning duration. These results suggest that mPFC VIP neurons play crucial roles in rapid reversal learning, but not in incremental cue-outcome association learning, by monitoring significant deviations from ongoing environmental contingency and imposing error-correction signals during behavioral adjustments. These findings shed light on the intricate cortical circuit dynamics underpinning behavioral flexibility in complex, multifaceted environments.

## INTRODUCTION

Behavioral flexibility is an essential skill for survival in an ever-changing world. We must adapt our behaviors flexibly in response to changes in environmental contingencies, which are often structured in a complex manner across multiple dimensions. While previous research has predominantly focused on the roles of specific brain regions in various aspects of behavioral flexibility^1-6^, a growing body of evidence indicates that processes supporting different types of behavioral flexibility are distributed across diverse brain areas^7-11^, highlighting the need to investigate how a given brain region integrates different forms of behavioral flexibility—a topic that has been largely overlooked.

We investigated this matter in the medial prefrontal cortex (mPFC), a key player in behavioral flexibility^12-15^, using mice performing a probabilistic reversal learning task. This task consists of two levels of environmental dynamics—stochastic outcome delivery and the reversal of cue-outcome contingencies—and thus tax two different types of learning: incremental value learning (or gradual cue-outcome association), which refines responses to the probabilistic nature of outcomes, and rapid reversal learning, which allows for rapid adaptation to sudden shifts in cue-outcome contingencies^16-19^. We have shown previously that variables related to incremental value learning are represented in widespread areas of the rodent brain including the mPFC^9,20^. We also have shown that mPFC inactivation significantly delays reversal learning in the current task^21^. Even though early studies using deterministic outcome delivery showed the importance of the orbitofrontal cortex in reversal learning and the mPFC in extradimensional set shifting in rodents^22-24^, we found that the mPFC inactivation strongly impairs probabilistic reversal learning^21^. These findings indicate that the mPFC is equipped with neural machineries for both incremental value learning and rapid reversal learning.

We focused on vasoactive intestinal polypeptide (VIP)-expressing neurons in the present study. Despite being a relatively small population, they are thought to exert powerful influences on cortical information processing because they control the activity of other cortical inhibitory interneurons^25-27^. Thus, changes in VIP neuronal activity are likely to be translated into global changes in cortical network activity. We therefore reasoned that VIP neurons may play a particularly important role in high flexibility-demanding situations, such as during reversal learning. To probe this hypothesis, we examined calcium dynamics and perturbation effects of mPFC VIP neurons while mice were performing the probabilistic reversal learning task. In particular, in order to obtain insights on mPFC neural circuit dynamics enabling rapid reversal learning, we focused on VIP neuronal activity dynamics while reversal learning is in progress, rather than merely comparing neural activity before and after reversal learning. We also analyzed fast-spiking (FS; putative inhibitory) and regular-spiking (putative pyramidal or PP) neuronal activities recorded in our previous study^21^ and compared them with VIP neuronal activity. The results indicate that VIP neurons are pivotal for reversal learning but not for gradual cue-outcome association learning.

## RESULTS

### Chemogenetic modulation of VIP neurons

To examine the effect of modulating VIP neuronal activity on probabilistic reversal learning, we bilaterally injected a Cre-dependent AAV carrying excitatory designer receptor exclusively activated by designer drug (DREADD) hM3D(Gq) (pAAV2-hSyn-DIO-hM3D(Gq)-mCherry) or inhibitory DREADD hM4D(Gi) (pAAV2-hSyn-DIO-hM4D(Gi)-mCherry) into the mPFC of VIP-Cre mice. After 4-9 weeks, we trained them in a probabilistic classical conditioning task under head fixation (**Fig. 1a-b;** see Methods). In this task (task 1), two different odor cues (conditioned stimulus or CS) were paired with a reward (2 ul of water) and a punishment (air puff delivery to the right eye; unconditioned stimulus or US) each with 75% probability, with a 1-s delay between the CS and the US. The mice were trained until their anticipatory lick rates differed significantly between reward-predicting cue (CS_rw_) and punishment-predicting cue (CS_pn_) trials (600 trials per session; total 1-3 sessions). They were then subjected to reversal learning, in which one randomly chosen reward-predicting cue was paired with air puff instead of water (CS_rw→pn_), and one randomly chosen punishment-predicting cue was paired with water instead of air puff (CS_pn→rw_) at 75% probability (**Fig. 2**). The mice were trained in reversal learning until they reached the reversal criterion in 300 trials within a single session (total 2-15 reversal sessions; see Methods for details).

**Fig. 1.**
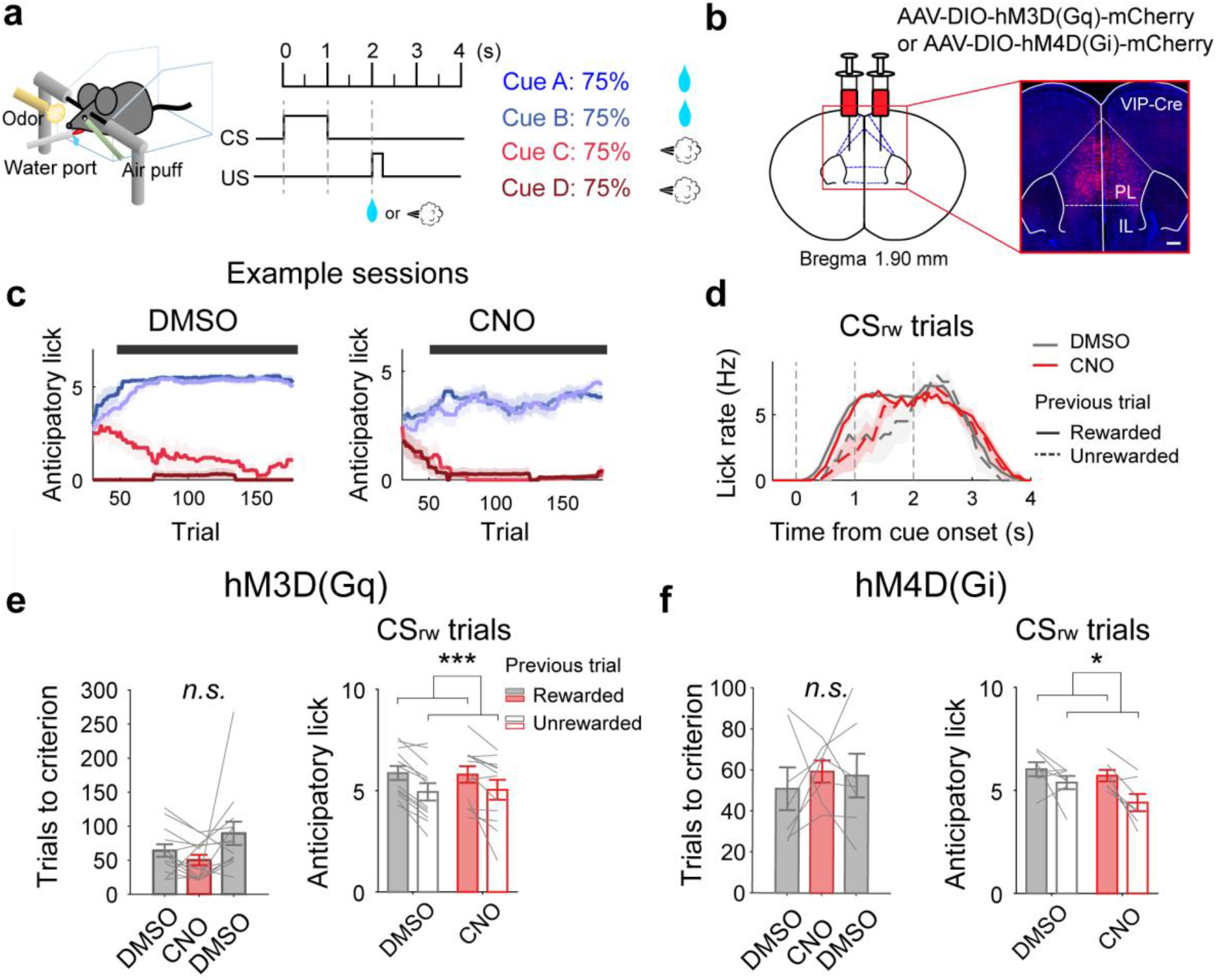
Effect of modulating VIP neuronal activity on probabilistic classical conditioning. **(a)** A schematic for task 1, used for chemogenetic and optogenetic experiments. Head-fixed mice performed a probabilistic classical conditioning task in which two odor cues (CS_rw_) were paired with water reward, while two others (CS_pn_) were paired with air puff (punishment) at 75% probability. **(b)** A schematic for viral vector injection and histological confirmation of hM3D(Gq) expression in the mPFC. Scale bar, 500 μm. **(c)** Sample DMSO (left) and CNO (right) sessions showing mean anticipatory lick numbers (1-s window since cue offset) across trials (moving average of 60 trials) during daily training until reaching performance criterion before reversal onset. The cue configuration was identical to that at the end of the previous session. Blue and red traces denote reward-predicting (CS_rw_) and punishment-predicting (CS_pn_) cues, respectively. Black squares on top denote significant differences in all CS_rw_ versus CS_pn_ comparisons (*p* < 0.05, Tukey-Kramer post hoc test following one-way ANOVA) in that trial. **(d)** Sample DMSO (red) and CNO (gray) sessions from an hM4D(Gi)-expressing mouse showing lick rates (500-ms moving average, 100-ms steps) grouped by reward delivery in the previous trial (solid and dashed lines; rewarded and unrewarded, respectively). Only consecutive CS_rw_ trials were analyzed. **(e and f)** Left, the number of trials to performance criterion (see Methods) with DMSO or CNO treatment. Right, anticipatory lick numbers grouped by drug and reward delivery in the previous trial for two consecutive CS_rw_ presentations. ^*^*p* < 0.05, ^***^*p* < 0.001, two-way repeated measures ANOVA. Gray lines, individual animal data. Shading (c and d), SEM across trials. Bar graphs and error bars (e and f), mean and SEM across animals (hM3D(Gq), n = 13; hM4D(Gi), n = 7).

**Fig. 2.**
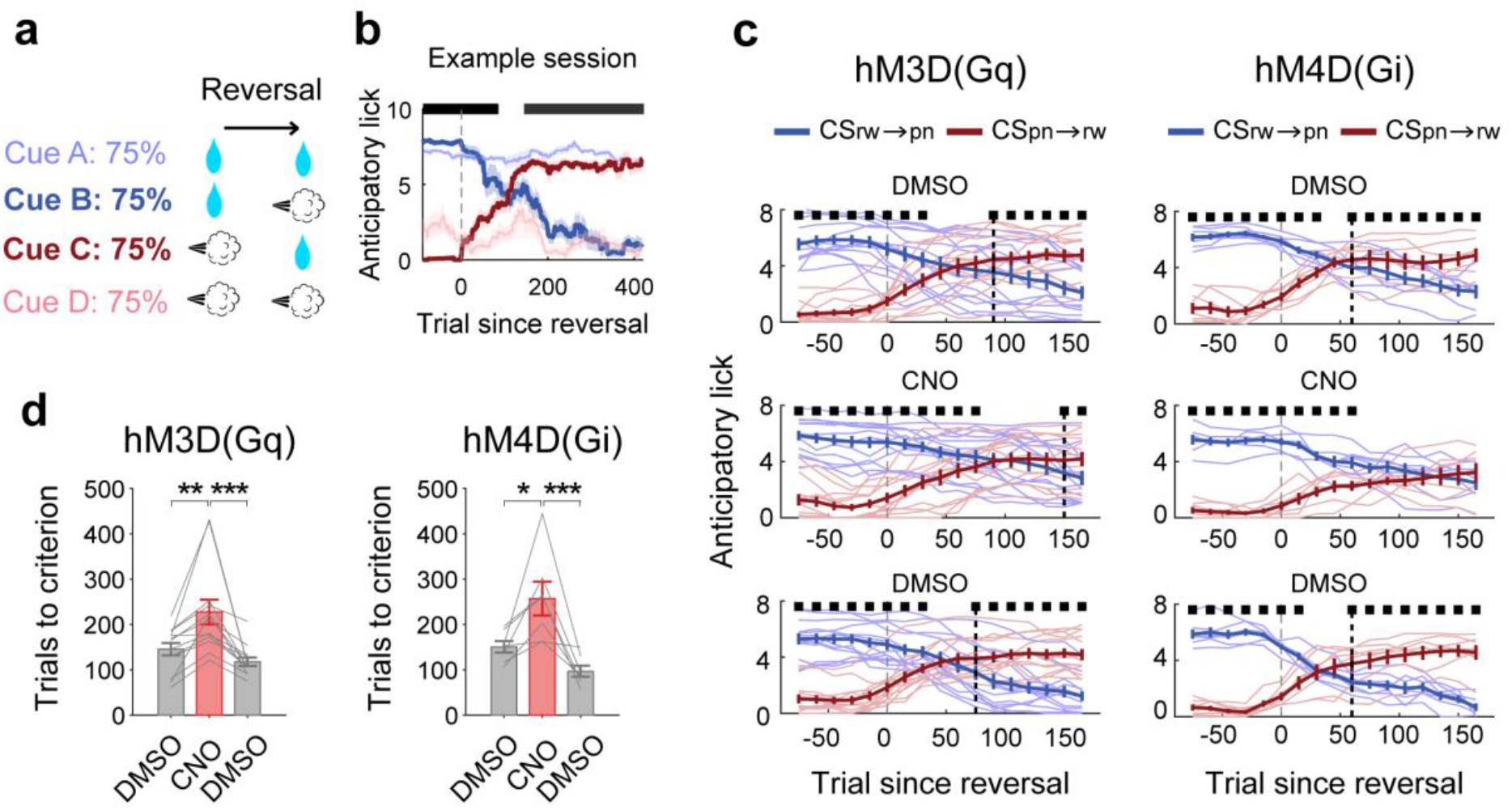
Effect of modulating VIP neuronal activity on reversal learning. **(a)** A schematic for reversal learning in task 1. One set of reward-predicting (CS_rw_) and punishment-predicting (CS_pn_) cues was subject to cue-outcome contingency reversal. **(b)** A sample session showing changes in cue-dependent anticipatory licking in the course of reversal learning. Saturated colors indicate the cue set subjected to reversal (blue, CS_rw→pn_; red, CS_pn→rw_) while soft colors denote the other cues (blue, CS_rw_; red, CS_pn_). The same format as in Fig. 1c. **(c and d)** Effects of chemogenetic activation (hM3D(Gq); left) or inactivation (hM4D(Gi); right) on reversal learning. **(c)** Anticipatory lick numbers in response to CS_rw→pn_ (blue) and CS_pn→rw_ (red) in the course of reversal learning (60-trial moving average, 15-trial steps). Thin lines, individual animal data. Thick lines, their averages. Black square, significant (*p* < 0.05, *t*-test) difference between two trial types (CS_rw→pn_versus CS_pn→rw_). Black dashed lines, first occurrence of significantly different anticipatory lick frequencies across animals between two trial types since reversal onset. **(d)** The number of trials to reversal criterion (see Methods) with DMSO or CNO treatment. Gray lines, individual animal data. Bar graphs, mean across animals. ^*^*p* < 0.05, ^**^*p* < 0.01, ^***^*p* < 0.001, Bonferroni post hoc test following one-way repeated measures ANOVA. Error bars and shading, SEM across trials (a) or animals (else) (hM3D(Gq), n = 13; hM4D(Gi), n = 7 mice).

We then tested the mice in daily reversal learning sessions with systemic injections (i.p.) of either dimethyl sulfoxide (DMSO; test sessions 1 and 3) or clozapine-N-oxide (CNO; test session 2). In each session, mice were initially trained in four-cue probabilistic conditioning until they met the performance criterion (see Methods) before reversal onset (**Fig. 1c**). Upon analyzing two consecutive CS_rw_ trials before reversal onset, we observed a significant increase in the number of anticipatory licks during the delay-period (1 s) in the second CS_rw_ trial when the mouse had been rewarded in the preceding trials, as opposed to when it was unrewarded, indicating that anticipatory licking is influenced by the history of past rewards (**Fig. 1d**). This observation aligns with the finding that animals and humans continuously update expected outcomes on a trial-by-trial basis in response to stochastic outcome delivery^9,28,29^.

Chemogenetic manipulation of VIP neurons did not significantly alter the number of trials to the performance criterion before reversal onset (one-way repeated measures ANOVA, hM3D(Gq): *F*(2,38) = 2.689, *p* = 0.082; hM4D(Gi): *F*(2,12) = 0.221, *p* = 0.805; **Fig. 1e-f left**), which is consistent with our previous finding that mPFC inactivation has minimal impact on probabilistic classical conditioning^21^. Chemogenetic manipulation of VIP neurons did not significantly affect the influence of past rewards on anticipatory licking, either (two-way repeated measures ANOVA, hM3D(Gq): main effect of previous reward, *F*(1,11) = 27.566, *p* = 2.7×10^-4^; drug, *F*(1,11) = 0.003, *p* = 0.955; previous reward×drug, *F*(1,11) = 0.405, *p* = 0.538; hM4D(Gi): main effect of previous reward, *F*(1,6) = 7.014, *p* = 0.038; drug, *F*(1,6) = 2.662, *p* = 0.154; previous reward×drug, *F*(1,6) = 2.893, *p* = 0.140; **Fig. 1e-f right**). Together, these results argue against the importance of mPFC VIP neurons in cue-outcome association learning before reversal onset.

In each daily reversal learning session, the stimulus-outcome pairing was reversed for only one pair of CS_rw_ and CSpn, with training continuing until the mouse reached the reversal criterion (see Methods; **Fig. 2a-b**). Chemogenetic activation of VIP neurons significantly increased the number of trials to the reversal criterion (hM3D(Gq): 13 mice; one-way repeated measures ANOVA, *F*(2,24) = 12.644, *p* = 1.8×10^−4^; Bonferroni’s post hoc test, 1^st^ DMSO vs. CNO, *p* = 0.002, 1^st^ DMSO vs. 2^nd^ DMSO, *p* = 0.600, CNO vs. 2^nd^ DMSO, *p* = 4.0×10^−4^; **Fig. 2c left, d left**). Similarly, chemogenetic inactivation of VIP neurons significantly increased the number of trials to the reversal criterion (hM4D(Gi): seven mice; *F*(2,12) = 13.009, *p* = 9.9 × 10^−4^; Bonferroni’s post hoc test, 1^st^ DMSO vs. CNO, *p* = 0.018, 1^st^ DMSO vs. 2^nd^ DMSO, *p* = 0.355, CNO vs. 2^nd^ DMSO, *p* = 5.0×10^−4^; **Fig. 2c right, d right**). These results indicate that a deviation from the normal level of mPFC VIP neuronal activity may impair probabilistic reversal learning.

### VIP neural correlates of classical conditioning

Next, we examined VIP neuronal activity during probabilistic reversal learning (**Fig. 3**). For this, we injected a Cre-dependent AAV carrying the calcium indicator jGCaMP7f (pGP-AAV1-syn-FLEX-jGCaMP7f-WPRE) unilaterally into the mPFC of VIP-Cre mice. Additionally, we implanted a GRIN prism lens targeting layers 2-3 which primarily harbor VIP neurons^27^ (**Fig. 3b**; see **Supplementary Fig. 1** for histological verification of GRIN prism lens locations). Following 8-22 weeks from surgery, the mice were trained in a modified version of the probabilistic classical conditioning task. In this task (task 2), three different odor cues were associated with 75% water delivery, 75% air puff delivery, and no outcome (CS_rw_, CSpn, and CSnt, respectively). Importantly, access to the water port for licking was restricted during cue presentation (1.5 s) and the first delay period (delay 1; 1.5 s; **Fig. 3a**). The water port was advanced toward the animal at delay 1 offset and, after an additional 1.5 s delay during which the animal could engage in licking behavior (delay 2), an outcome was disclosed (**Fig. 3a**). This was to examine cue-dependent VIP neuronal activity with minimal interference from motor-related activity during the cue and initial delay periods, while also assessing cue-dependent anticipatory licking responses during the second delay period (**Fig. 3a**). After being trained in the task to the performance criterion (see Methods; 1-3 daily sessions; **Fig. 3a**), calcium imaging was performed (**Fig. 3b**).

**Fig. 3.**
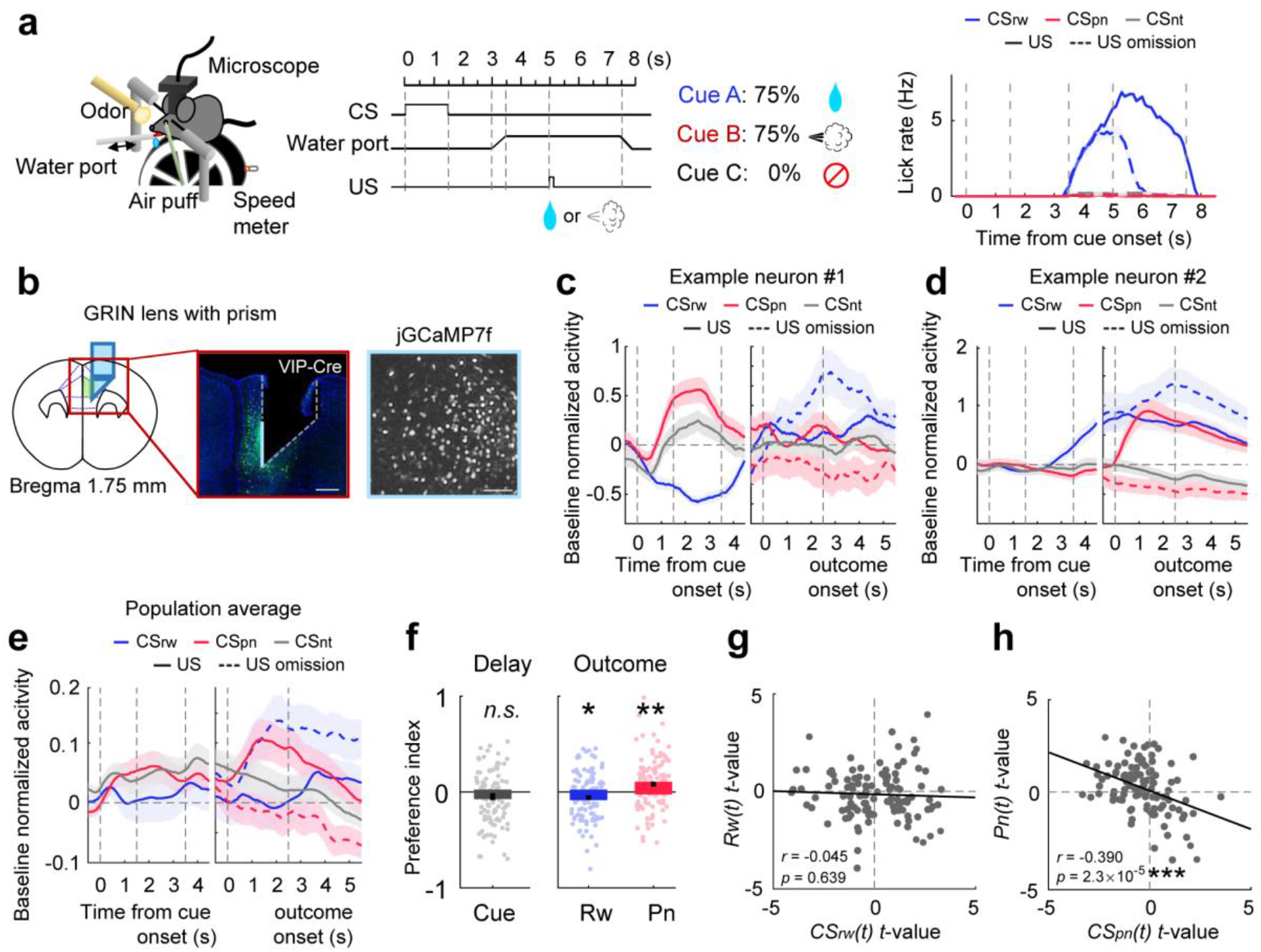
VIP neuronal activity during probabilistic classical conditioning. **(a)** A schematic for task 2, used for calcium imaging. Three odor cues were paired with a reward (75%; CS_rw_), an air puff (75%; CS_pn_), or no outcome (100%; CSnt). The water port was advanced toward the animal 1.5 s after cue offset and then retracted 4.5 s later. Right, time courses of cue-dependent lick rate during a sample session. (**b)** A schematic (left) and a coronal brain section (middle) showing the GRIN prism lens position and the spread of jGCaMP7f (green) in the mPFC. Right, a sample field of view. Scale bar, middle, 500 μm; right, 100 μm. **(c)** Cue-dependent responses of a sample VIP neuron. **(d)** Outcome-dependent responses of a sample VIP neuron. **(e)** Mean baseline-normalized population responses of VIP neurons. **(f)** Left, mean cue preference index (+ and -, higher activity in response to CS_rw_, and CS_pn_, respectively, see Methods) during the first delay period (1.5 s since cue offset). Right, mean outcome-preference indices (+ and -, higher activity in response to US-delivery and US-omission, respectively, see Methods) for reward (Rw) and punishment (Pn) during the outcome period (1.5 s since outcome onset). ^*^*p* < 0.05, ^**^*p* < 0.01, difference from zero, *t*-test. **(g and h)** Outcome-period VIP neuronal activity was analyzed using multiple linear regression to relate *t*-values of reward-predicting cue (*CS*_*rw*_*(t)*; eq. 3) and reward (*Rw(t)*; eq. 1; **g**) as well as *t*-values of punishment-predicting cue (*CS*_*pn*_*(t)*; eq. 3) and punishment (*Pn(t)*; eq. 2; **h**). Each circle represents one VIP neuron. *r*, Pearson’s correlation coefficient. The lines are results of linear regression. Error bars and shading, SEM across trials (c and d) or neurons (e to h; 111 neurons recorded from nine animals).

We first explored VIP neuronal correlates of classical conditioning. For each mouse, we analyzed the session after the mouse meeting the performance criterion for task 2, just before beginning reversal training. In this analysis, we included only those neurons with calcium event rates during task performance ≥ 0.025 Hz (111 VIP neurons from nine mice). Fig. 3c and 3d show sample VIP neuronal responses, and Fig. 3e shows the mean population responses. Throughout the study, unless noted otherwise, neural responses are shown as z-normalized activity using the mean and SD of neural activity during 0.5 s before cue onset (baseline normalization). As shown, individual VIP neurons showed diverse arrays of cue-dependent and outcome-dependent activity (**Fig. 3c-e**). The cue-preference index (positive and negative values, stronger responses to reward-predicting and punishment-predicting cues, respectively; see Methods) during the first delay period did not differ significantly from zero (*t*-test, *t*_*110*_ = -1.345, *p* = 0.181), indicating that VIP neuronal responses are not biased toward the reward-predicting or punishment-predicting cue (**Fig. 3f left**). During the outcome period, VIP population activity was lower in rewarded than unrewarded trials following CS_rw_ presentation and, conversely, higher in punished than unpunished trials following CSpn presentation. Consequently, the outcome-preference index (positive and negative values, stronger responses to US-delivery and US-omission, respectively; see Methods) was significantly negative for reward (*t*_*110*_ = -2.530, *p* = 0.013) and positive for punishment (*t*_*110*_ = 3.270, *p* = 0.001; **Fig. 3f right**). Additionally, VIP neurons were significantly responsive to diverse events during the task, such as CS onset, so that VIP neuronal population activity carried precise information about the progress of a trial (**Supplementary Fig. 2**). These results show that mPFC VIP neurons carry a diverse array of task-related information.

To investigate VIP neuronal activity related to outcome evaluation and value updating, we performed multiple regression analyses (eq. 1-3) and examined how cue-related and outcome-related responses during the outcome period (the first 1.5 s since outcome onset) are related. We especially focused on potential error-correction signals related to expected reward and punishment. Given that reward prediction error (RPE) is the difference between the expected and actual rewards, an RPE-coding neuron is expected to show responses to the reward-predicting cue and reward (*CS*_*rw*_*(t)*) and *Rw(t)* terms in multiple regression, respectively) in opposite directions^30^. Furthermore, if a neural population is predominantly involved in RPE representation, it should manifest a negative relationship between *CS*_*rw*_*(t)*- and *Rw(t)*-related activity. We found no significant negative relationship between the *t*-values of *CS*_*rw*_*(t)* and *Rw(t)* (Pearson’s correlation, *r* = -0.045, *p* = 0.639, **Fig. 3g**). However, a significant negative correlation was observed between the *t*-values of punishment-predicting cue (*CS*_*pn*_*(t)*) and punishment (*Pn(t)*; *r* = -0.390, *p* = 2.3×10^−5^, **Fig. 3h**). These results suggest that mPFC VIP neurons are involved in updating expected punishment but not expected reward during probabilistic classical conditioning.

### VIP neuronal activity before and after reversal

Following initial training in task 2, mice were further trained in reversal learning until reaching the reversal criterion in ≤ 150 trials (total 2-7 daily sessions; **Fig. 4a**). In this phase, after cue-outcome contingency reversal, the CS_rw_ was paired with 75% air puff instead of water (CS_rw→pn_) and the CS_pn_ with 75% water instead of air puff (CS_pn→rw_). We then recorded VIP neuronal activity during daily reversal learning sessions, with one episode of reversal occurring per session (reversal onset trial, 141-160; trials to reversal criterion, 46-130; total, 420 trials). One reversal session from each mouse was chosen for analysis based on behavioral performance and recording quality, and only neurons with calcium event rates during task performance ≥ 0.025 Hz were included in the analysis (106 VIP neurons recorded from 12 mice).

**Fig. 4.**
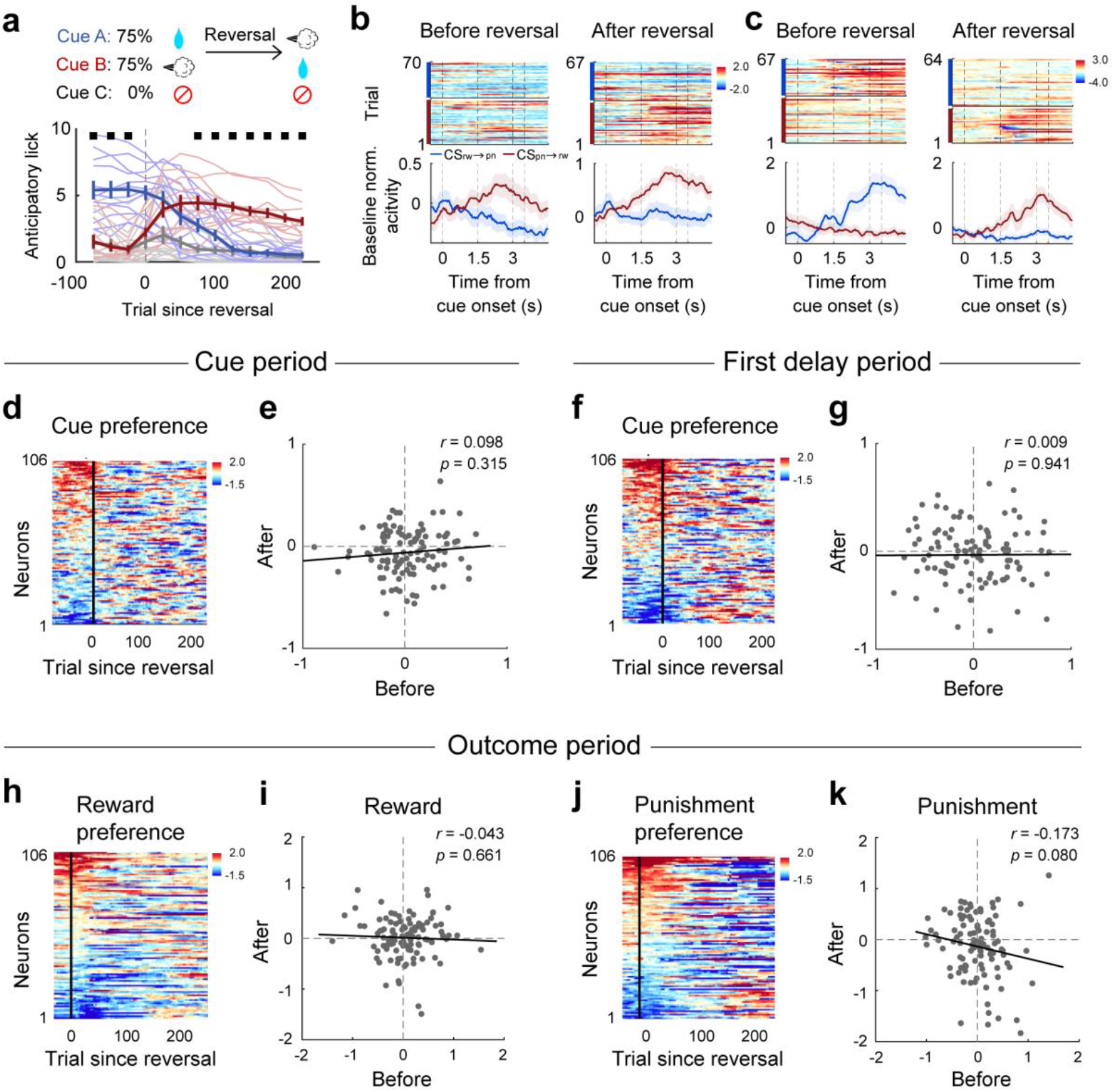
VIP neuronal activity before and after reversal. **(a)** Top, a schematic for reversal learning in task 2. Bottom, behavioral performance of all mice used for VIP neuronal recordings. Shown are cuedependent anticipatory lick numbers during the second delay period in the course of reversal learning (moving average of 50-trials, 25-trial steps). The same format as in Fig. 2c. **(b and c)** Cue-dependent responses of sample VIP neurons before (left) and after (right) reversal. Upper, heat maps of baseline-normalized fluorescence; Lower, their averages. Trials were grouped by odor cues (CS_rw→pn_(blue) and CS_pn→rw_ (red)). **(d)** Dynamics of cue-preference index (40-trial moving average; color-coded) during the cue period (1.5 s since cue onset) in the course of reversal learning. Neurons were aligned according to their cue-preference index before reversal onset. **(e)** Cue-preference index during the cue period before (100 trials before reversal onset; abscissa) and after (151-250 trials since reversal onset; ordinate) reversal. The same format as in Fig. 3g. **(f and g)** Dynamics of cue-preference index during the first delay period (1.5 s since cue offset) in the course of reversal learning. The same format as in (d and e). **(h to k)** Dynamics of reward (h and i) and punishment-preference indices (j and k) during the outcome period (1.5 s since outcome onset). The same format as in (d and e). Error bars and shading, SEM across animals (a) or trials (b and c) (n = 106 neurons recorded from 12 animals).

Across reversal, VIP neurons showed a diverse array of activity patterns (**Fig. 4b-c**). We have shown previously that the majority of mPFC neurons reverse their cue-preference index across reversal so that they are consistently responsive to expected outcomes rather than cue identity^21^. However, VIP neurons showed mixed response changes: some responded consistently to cue identity before and after reversal **(Fig. 4b)**, while others responded primarily to expected outcomes **(Fig. 4c)**. Additionally, some VIP neurons did not show consistent responses to neither cue identity nor expected outcomes, resulting in no significant relationship in the cue-preference index before and after reversal (**Fig. 4d-g**). Moreover, VIP neurons exhibited mixed response changes to both reward and punishment before and after reversal (**Fig. 4h-k**). These findings reveal that the activity of the VIP neuronal population undergoes dynamic and complex changes during reversal, rather than exhibiting simple sensory or expected outcome-dependent activity.

### VIP neuronal responses to reversal onset

To investigate how VIP neurons contribute to reversal learning, we first examined whether VIP neurons contribute to the detection of reversal onset. Specifically, we examined whether the cue and outcome responses of the VIP neuronal population undergo sudden activity changes after encountering an unexpected outcome. For this analysis, neuronal activity was normalized to that during 100 trials before reversal onset (trial normalization; see Methods). Remarkably, upon the mouse experiencing the first unexpected outcome since reversal onset, the VIP neuronal population showed an abrupt and transient cue response change; the first cue response following the first unexpected outcome decreased significantly from the pre-reversal baseline level (*t*-test, *t105* = -3.012, *p* = 0.003), and then gradually returned to the baseline level within four appearances of the same cue (**Fig. 5a-b**). We failed to find such cue response dynamics when we analyzed PP and FS neurons that were recorded in a separate study^21^ (**Supplementary Fig. 3**). Upon separately examining VIP neuronal populations depending on whether the first unexpected outcome was a reward or punishment, we found the abrupt and transient cue response change in the reward-first group (baseline versus the first cue response since unexpected outcome, 69 VIP neurons recorded from six mice, *t*_*68*_ = - 3.736, *p* = 3.9×10^-4^), but not in the punishment-first group (37 VIP neurons recorded from six mice, *t*_*36*_ = -0.341, *p* = 0.735; **Fig. 5c**). These results indicate that VIP neuronal population signal a significant deviation in ongoing environmental contingency by abruptly and transiently decreasing their cue-related activity following an unexpected reward encounter since reversal onset.

**Fig. 5.**
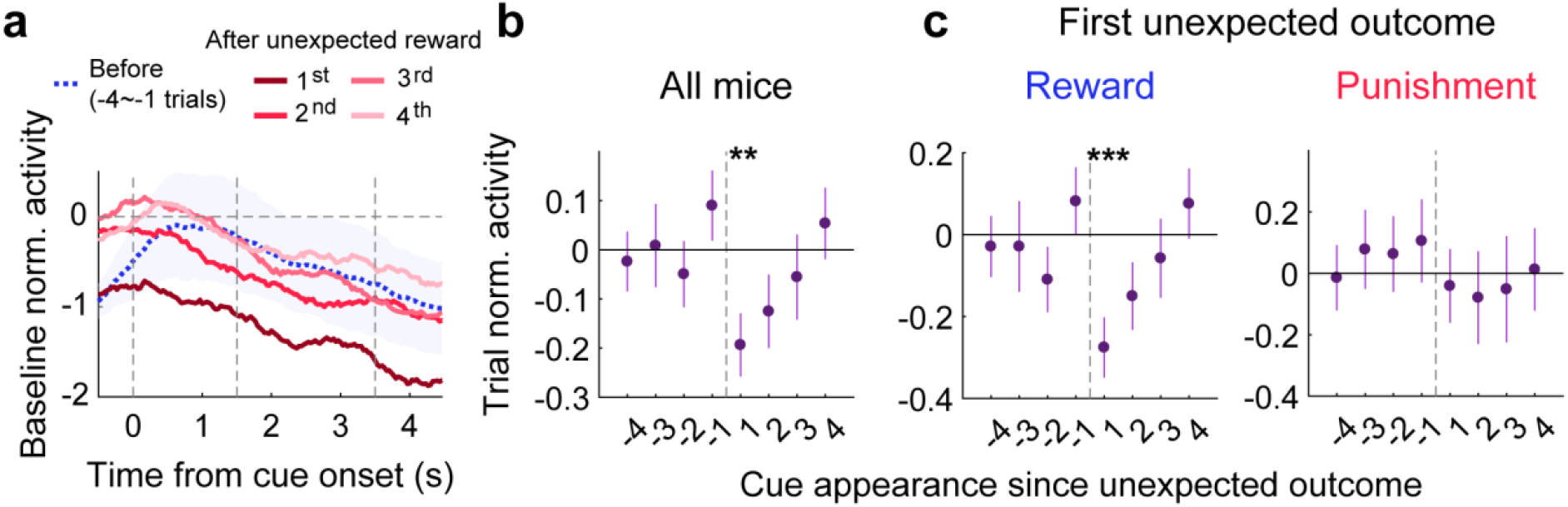
VIP neuronal responses to reversal onset. **(a)** Baseline-normalized individual-trial responses of a sample VIP neuron to CS_rw→pn_before (4-trial average) and after the first unexpected reward delivery since reversal onset. **(b)** Group data showing mean (±SEM across 106 neurons recorded from 12 mice) cue responses of VIP neurons during the first delay period, normalized to pre-reversal responses (100 trials; trial normalization), before and after experiencing the first unexpected outcome delivery. The abscissa denotes the order of the appearance of a given cue (CS_rw→pn_or CS_pn→rw_) around the first unexpected outcome delivery. Cue responses are shown as the mean of CS_rw→pn_ and CS_pn→rw_ responses. **(c)** The same format as in (b), but sessions were divided according to the type of the first unexpected outcome (reward versus punishment; 69 neurons from six animals and 37 neurons from six animals, respectively) since reversal onset. ^**^*p* < 0.01, ^***^*p* < 0.001, difference from zero, *t*-test.

The dynamics of outcome responses since reversal onset differed from those of cue responses (**Supplementary Fig. 4**). VIP neuronal population responses to the first encounter of an unexpected reward or punishment did not deviate significantly from the baseline. Instead, their responses to reward and punishment changed significantly from the baseline activity only upon the second encounter of the unexpected outcome (**Supplementary Fig. 4b-c**), which can be attributed to the altered cue responses of VIP neuronal population since reversal onset. This suggests that VIP neuronal population does not respond to an unexpected reward delivery per se, but integrates this information into its cue responses in the subsequent trial.

### Error-correction signals during reversal learning

We then investigated whether VIP neurons play a role in revising cue-associated expected outcomes during reversal. Using multiple regression analyses (eq. 1-3), we examined how cue and outcome responses during the outcome period (first 1.5 s) are related in the course of reversal learning. Specifically, we examined the relationships between responses to the reward-predicting cue (*CS*_*rw*_*(t)*; CS_rw→pn_and CS_pn→rw_ before and after reversal onset, respectively) and reward in the current trial (*Rw(t)*), and between responses to the punishment-predicting cue (*CS*_*pn*_*(t)*; CS_pn→rw_ and CS_rw→pn_before and after reversal onset, respectively) and punishment in the current trial (*Pn(t)*). We found a significant negative relationship between *t*-values of *CS*_*rw*_*(t)* and *Rw(t)* during the reversal phase (1-100 trials since reversal onset; Pearson’s correlation, *r* = -0.378; *p* = 6.3×10^-5^), but not before (1-100 trials before reversal onset; *r* = 0.182; *p* = 0.062; also see **Fig. 3g**) or after (151-250 trials since reversal onset; *r* = -0.009; *p* = 0.925; **Fig. 6a**). This suggests the engagement of VIP neurons in conveying RPE signals during reversal. In comparison, analysis of PP and FS neuronal activity from our previous study^21^ revealed different patterns (eq. 4-6; **Supplementary Fig. 5**). The PP neuronal population showed a significant negative relationship between the *t*-values of *CS(t)* and *Rw(t)* in all three phases (before, during and after reversal; *r* = -0.174, -0.227 and -0.210; *p* = 0.002, 5.2×10^-5^ and 1.9×10^-4^, respectively; **Supplementary Fig. 5a**), suggesting their routine involvement in conveying RPE signals irrespective of cue-outcome contingency reversal. In contrast, the FS neuronal population showed no significant relationship between *t*-values of *CS(t)* and *Rw(t)* in all three phases (before, during and after reversal; *r* = 0.170, - 0.025 and -0.047; *p* = 0.169, 0.840 and 0.707, respectively; **Supplementary Fig. 5e**). These results indicate that reversal phase-specific engagement in RPE coding is a prominent feature of VIP neuronal population.

**Fig. 6.**
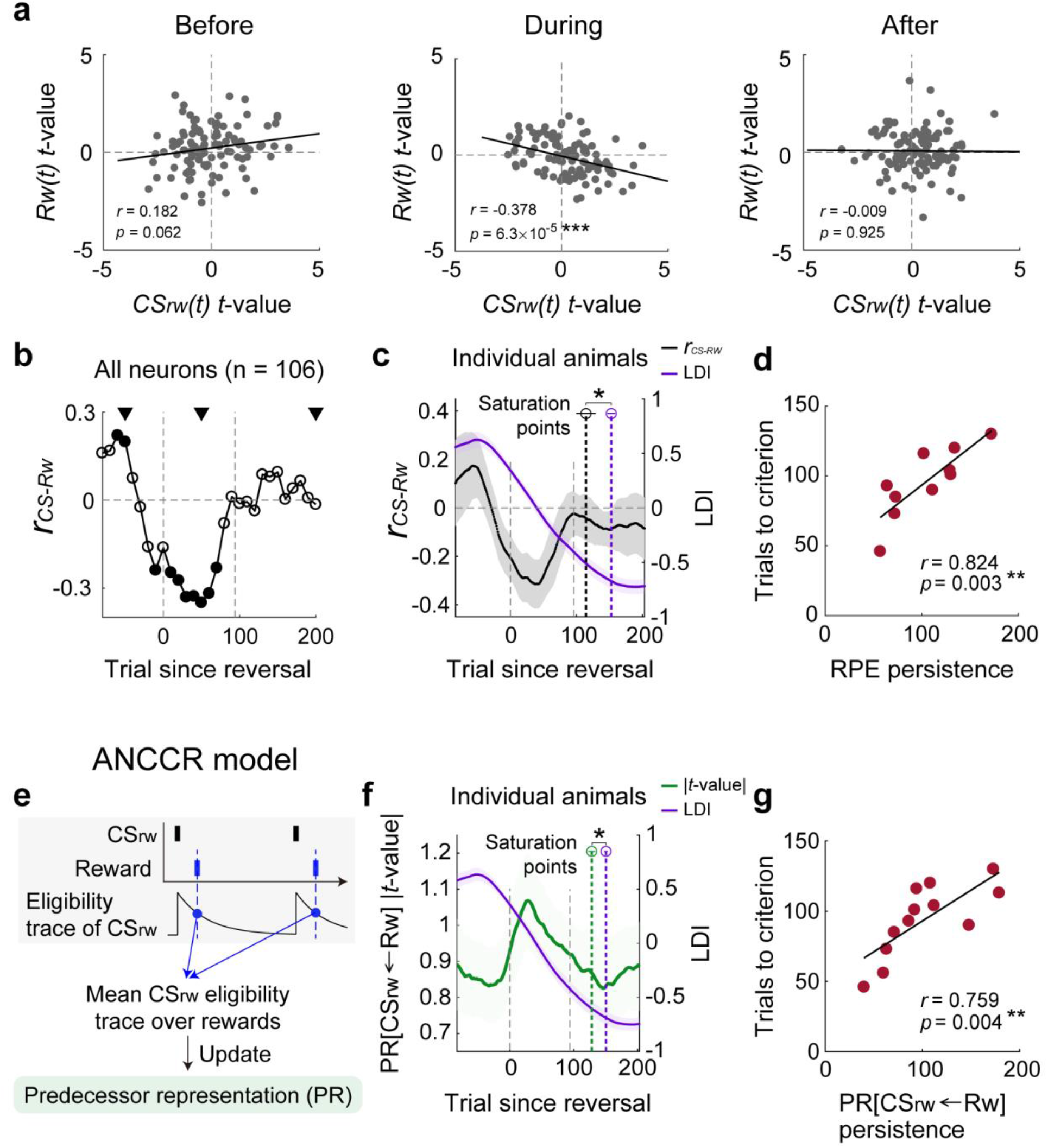
Error correction signals during reversal learning. **(a to d)** Outcome-period VIP neuronal responses to reward-predicting cue (*CS*_*rw*_*(t)*; CS_rw→pn_and CS_pn→rw_ before and after reversal, respectively) and reward (*Rw(t)*) were analyzed using multiple linear regression as in Fig. 3g-h. **(a)** The scatter plots show *t*-values for *CS*_*rw*_*(t)* (abscissa) and *Rw(t)* (ordinate) before, during, and after reversal (left, middle, and right black triangles in b, respectively) for each neuron (n = 106 neurons recorded from 12 animals). The same format as in Fig. 3g. **(b)** Pearson’s correlation coefficient between *t*-values for *CS*_*rw*_*(t)* and *Rw(t)* (*r*_*CS-Rw*_) of all VIP neurons during reversal (100-trial moving window, 10-trial steps). Left and right dashed vertical lines, reversal onset and the mean number of trials until reversal criterion, respectively. Filled circles, *p* < 0.05 (difference from 0). **(c)** Mean temporal profiles of LDI (+ and -, higher anticipatory lick number in response to CSrw→pn, and CS_pn→rw_ presentation, respectively, see Methods; purple line) and smoothed *r*_*CS-Rw*_ (100-trial moving window, 1-trial steps, 40-trial smoothing; black line) across animals (n = 10) during reversal. Black and purple open circles and dashed vertical lines denote the mean saturation points for *r*_*CS-Rw*_ and LDI, respectively. Dashed vertical gray lines, reversal onset and the mean number of trials until reversal criterion. ^*^*p* < 0.05, *t*-test. **(d)** The relationship between the persistence of RPE signals (the number of trials between the minimum and the maximum *r*_*CS-Rw*_-values after reversal onset; abscissa) and the number of trials to reversal criterion (ordinate). Circles, individual animal data. **(e)** A schematic for calculating predecessor representation of CS_rw_ over reward (PR[CS_rw_ ←Rw]) in the ANCCR model. **(f)** Mean temporal profiles (n = 12 mice) of LDI (purple line) and absolute *t*-value for *PR[CS*_*rw*_ *←Rw](t)* (green line) obtained from multiple linear regression (eq. 9) of outcome-period VIP neuronal responses (100-trial moving window, 1-trial steps, 40-trial smoothing) during reversal. Green and purple open circles and dashed vertical lines denote the mean saturation points for *PR[CS*_*rw*_ *←Rw](t)* absolute *t*-value and LDI, respectively. Dashed vertical gray lines, reversal onset and the mean number of trials until reversal criterion. ^*^*p* < 0.05, *t*-test. **(g)** The relationship between the persistence of PR[CS_rw_ ←Rw] signals (the number of trials between the maximum and minimum absolute *t*-values for *PR[CS*_*rw*_ *←Rw](t)* after reversal onset; abscissa) and the number of trials to reversal criterion (ordinate). Circles, individual animal data. ^*^*p* < 0.05, ^**^*p* < 0.01, ^***^*p* < 0.001. Error bars and shading, SEM across animals.

Regarding punishment, all three neuronal populations showed a significant negative relationship between *t*-values of *CS*_*pn*_*(t)* and *Pn(t)* in all phases of reversal (before, during and after reversal, VIP neurons, *r* = -0.420, -0.501 and -0.591; *p* = 7.5×10^-6^, 4.4×10^-8^ and 2.6×10^-11^, respectively; PP neurons, *r* = -0.301, -0.511 and -0.467; *p* = 6.0×10^-8^, 3.5×10^-22^ and 2.6×10^-18^, respectively; FS neurons, *r* = -0.389, -0.280 and -0.545; *p* = 0.001, 0.022, and 1.8×10^-6^, respectively; **Supplementary Fig. 6a, 5c, g**). This indicates that all three types of neurons routinely transmit punishment prediction error (PPE) signals before and after reversal (**Supplementary Fig. 5c-d, g-h, 6**). Thus, unlike the pattern observed with reward prediction, VIP neuronal population does not play a phase-specific role in conveying signals related to punishment prediction.

### Temporal dynamics of error-correction signals

To explore temporal dynamics of RPE signals during reversal, we analyzed the Pearson’s correlation between *t*-values of *CS*_*rw*_*(t)* and *Rw(t)* (*r*_*CS-Rw*_) in a moving window of 100 trials (in 10-trial steps). Upon the onset of reversal, *r*_*CS-Rw*_ rapidly declined, reaching its minimum at approximately 50 trials since reversal onset. It then increased gradually, approaching 0 as the animal’s licking behavior met the reversal learning criterion (**Fig. 6b**). This distinctive pattern was not observed in PP and FS neurons (**Supplementary Fig. 5b, f**). These results further support reversal phase-specific RPE coding as a distinctive feature of VIP neuronal population.

To further elucidate the relationship between the dynamics of VIP neuronal RPE signals and reversal learning, we examined the relationship between the rate of RPE signal change during reversal and the speed of reversal learning. For the former, we smoothed the temporal profile of each animal’s *r*_*CS-Rw*_ (100-trial window, 1-trial step) by employing a 40-trial moving average to mitigate the impact of noisy fluctuations, and calculated the difference in trials between the maximum *r*_*CS-Rw*_ after the animal reached the reversal criterion and the minimum *r*_*CS-Rw*_ within 93 trials since reversal onset (the average number of trials to reversal criterion; **Fig. 6c**). Concerning the latter, we used the number of trials to reversal criterion. Remarkably, our analysis revealed a strong positive correlation between these two measures (Pearson’s correlation, *r* = 0.824, *p* = 0.003; **Fig. 6d**). These findings were consistent across various standards for calculating the rate of RPE change (**Supplementary Fig. 7a-c**), and after excluding early analysis windows including pre-reversal trials (**Supplementary Fig. 7a, d-e**).

**Fig. 7.**
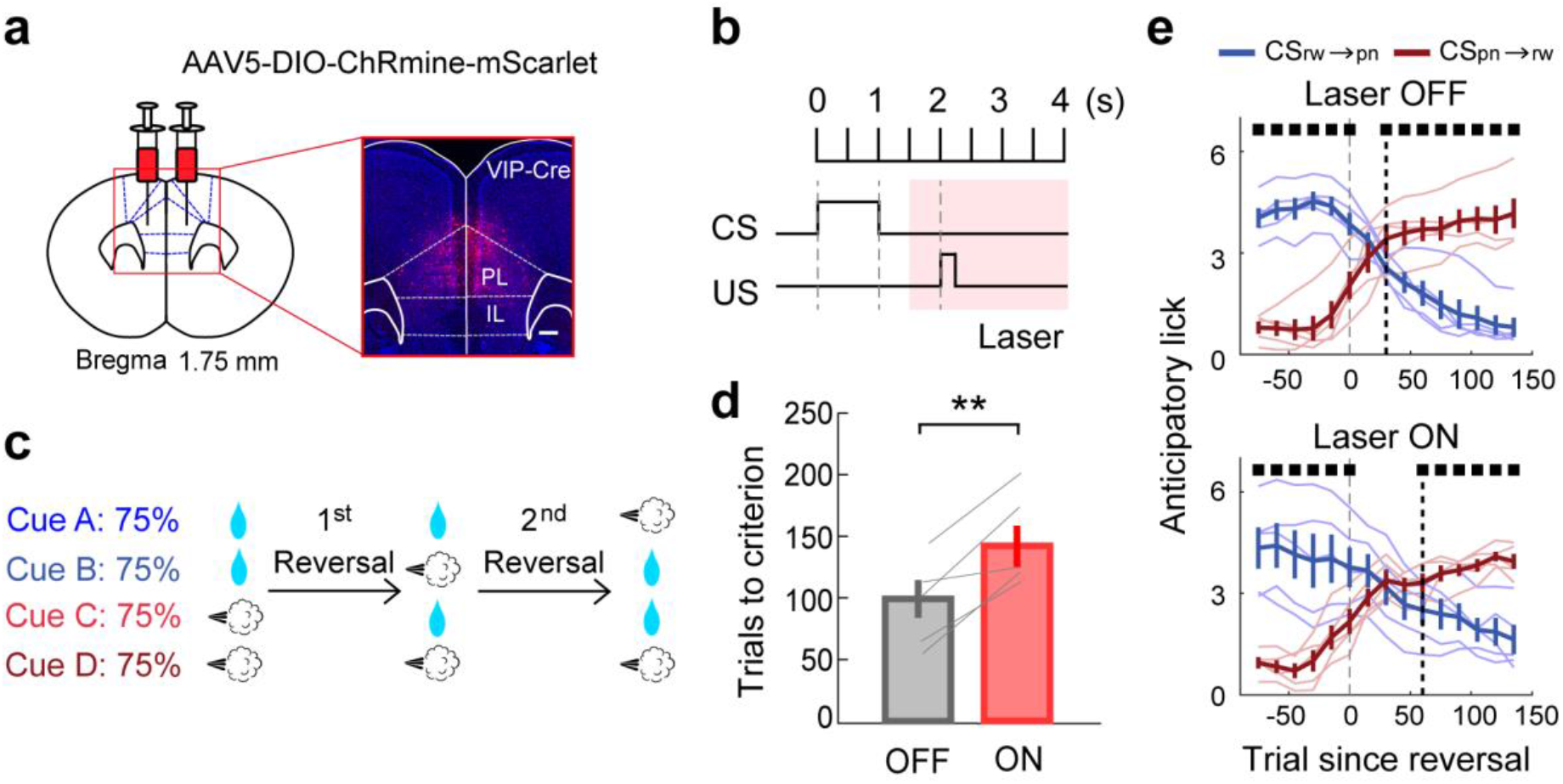
Effect of modulating outcome-period VIP neuronal activity on reversal learning. **(a)** A schematic for viral vector injection and histological confirmation of ChRmine expression in the mPFC. Scale bar, 500 μm. **(b)** Laser stimulation (633 nm, 20 Hz, 5-ms pulse; pink shading) was given during the outcome period in task 1. **(c)** A schematic for two reversals per session (task 1). Laser stimulation was applied only during one of two reversals. **(d)** The number of trials to reversal criterion with (red) or without (gray) laser stimulation. Gray lines, individual animal data. ^**^*p* < 0.01, *t*-test. **(e)** Changes in anticipatory lick frequency during reversal (60-trial moving average, 15-trial step). The same format as in Fig. 2c. Error bars, SEM across animals (n = 5).

To compare the time courses of RPE-related VIP neuronal activity dynamics with reversal learning, we determined the saturation points of the smoothed *r*_*CS-Rw*_ and the lick difference index (LDI, an index for cue-dependent anticipatory licking; see Methods). We found that the RPE saturation preceded the LDI saturation (defined as when an index passes the 95% of the difference between maximum and minimum values) by approximately 40 trials (38.40±16.78 trials; 112.60±14.57 and 151.00±5.85 trials since reversal onset, respectively; *t*-test, *t9* = -2.289, *p* = 0.048; **Fig. 6c**), suggesting that the dynamics of VIP neuronal RPE signals precede reversal learning. This finding is consistent with a causal role of RPE-related VIP neuronal activity in reversal learning.

Given the recent proposal that midbrain dopamine neurons may transmit retrospective association signals rather than traditional RPE signals^31^, we analyzed our data using the proposed ‘adjusted net contingency of causal relation’ (ANCCR) model. Specifically, we computed the retrospective association (predecessor representation or PR) of CS_rw_ over reward (PR[CS_rw_←Rw]) through a trial-by-trial analysis without employing a sliding window (details in ^31^; **Fig. 6e**). The absolute *t*-value for *PR[CS*_*rw*_*←Rw](t)* (eq. 9) showed an increase upon reversal onset followed by a subsequent decrease (**Fig. 6f**), mirroring the dynamics seen in *r*_*CS-Rw*_ (**Fig. 6c**). The difference in trials between the maximum absolute *t*-value (within the average number of trials to reversal criterion) and the minimum value (after the animal reached the reversal criterion) showed a significant correlation with the speed of reversal learning (Pearson’s correlation, *r* = 0.759, *p* = 0.004; **Fig. 6g**). Furthermore, the saturation of absolute *t*-value for *PR[CS*_*rw*_*←Rw](t)* also preceded the LDI saturation significantly (*t*-test, *t11* = -2.688, *p* = 0.021, **Fig. 6f**). These results provide additional support for the relationship between VIP neuronal activity dynamics and reversal learning.

### Optogenetic modulation of VIP neurons

The aforementioned findings underscore the importance of VIP neuronal RPE signals during the outcome period in reversal learning. If so, modulating VIP neuronal activity selectively during the outcome period should have an impact on reversal learning. To test this prediction, we bilaterally injected a Cre-dependent adeno-associated virus (AAV5-DIO-ChRmine-mScarlet), carrying the gene for highly sensitive red-shifted opsin ChRmine for transcranial activation, into the mPFC of VIP-Cre mice (n = 5, **Fig. 7a**), and attached an optic fiber to the skull. After a 4 to 7-week period following virus injection, during which the animals were trained to perform two episodes of reversal learning in each daily session (task 1; **Fig. 7c**), we applied light stimulation (intensity, 800 mW mm^-2^; pulse duration, 5 ms; frequency, 20 Hz) specifically during the outcome period (2.5 s; **Fig. 7b**). The light stimulation was given during either the first or the second episode of reversal learning alternately across daily sessions. This manipulation led to a significant increase in the number of trials required to reach the reversal criterion in the experimental group (*t*-test, *t4* = 4.827, *p* = 0.009; **Fig. 7d**), but not in the control group injected with the tdTomato-expressing virus lacking opsin (*t4* = -0.100, *p* = 0.925; **Supplementary Fig. 8**). These results confirm that selectively manipulating VIP neuronal activity during the outcome period alone is sufficient to delay reversal learning (**Fig. 7e**).

## Discussion

To obtain insights into the mechanisms by which the mPFC manages diverse forms of behavioral flexibility, we examined the role of VIP neurons in probabilistic reversal learning. Our results support the hypothesis that VIP neurons are pivotal in rapid adjustment of behavior in high flexibility-demanding situations, but not in routine trial-by-trial adjustments of behavior. First, perturbing VIP neuronal activity had no significant effect on pre-reversal performance or past outcome-dependent anticipatory licking behavior. However, it significantly delayed reversal learning. Second, VIP neurons showed abrupt and transient CS-related activity changes after encountering the first unexpected reward since reversal onset. Third, VIP neurons transmitted RPE signals during behavioral reversal, but not before and after, with the persistence of RPE signals negatively correlated with the speed of reversal learning. This is in contrast to PP neuronal activity, which conveyed trial-by-trial RPE signals throughout all phases of reversal learning. Collectively, these findings consistently indicate that mPFC VIP neurons play an important role in reversal learning, but make little contribution to the routine process of trial-by-trial value updating in an uncertain environment. Behavioral studies have generally focused on the roles of distinct brain regions in varying types of flexibility, while physiological studies have tended to focus on one form of flexibility at a time. Consequently, the mechanisms by which a particular neural circuit manages diverse forms of behavioral flexibility in complex environments have remained unclear. Our study shows that a specific cortical neuron type plays a selective role in a specific type of behavioral flexibility, showcasing the possibility of dissecting the detailed cortical circuit processes that enable optimal behavioral adjustment to complex environmental dynamics.

Early studies demonstrated strong VIP neuronal responses to aversive stimuli^32,33^. However, in our study, punishment-related responses subsided rapidly after initial exposure (**Supplementary Fig. 9**). Moreover, the engagement of VIP neurons in conveying error correction signals during the reversal phase was specifically tied to reward prediction, whereas they transmitted punishment-related error correction signals indiscriminately across all phases of reversal learning. Additionally, abrupt changes in VIP neuronal activity since reversal onset were triggered by unexpected reward delivery but not by unexpected punishment delivery. These findings suggest that mPFC VIP neurons may facilitate rapid reversal learning predominantly through the processing of reward, rather than punishment. Although the underlying source of this asymmetry remains unclear, previous research in monkeys has demonstrated preferential processing of neural signals associated with rewarded actions over unrewarded ones. This bias may reflect the evolutionary importance of remembering contexts and actions associated with rewards for survival in natural environments^34^.

We analyzed reward-related VIP neuronal activity through the lens of RPE as well as the ANCCR model. RPE-based learning models posit that learning commences with the prospective prediction of future outcomes from a current event. Conversely, the ANCCR model hypothesizes that learning starts by retrospectively inferring the causes of meaningful outcomes such as rewards^31^. In the ANCCR model, PR represents the retrospective association between CS_rw_ and reward, which is updated with the mean CS_rw_ eligibility trace over rewards. We found that VIP neuronal signals for the core components of the two learning paradigms—RPE and PR—become stronger transiently after reversal onset. Remarkably, the persistence of both signals correlated significantly with behavioral performance (**Fig. 6b-g**). Even though our results do not provide insights into which of the two models represents the psychological processes underlying the current probabilistic reversal learning more accurately, consistent findings from the analyses using both models corroborate the conclusion that VIP neurons play a significant role in facilitating rapid reversal learning through their unique activity during, but not before or after, reversal learning.

In our earlier work, we demonstrated that mPFC PV neurons display opposite responses to reward and punishment and, based on simple neural network modeling, proposed that such divergent responses may expedite probabilistic reversal learning^21^. Together with the current findings, this suggests a potential division of labor, with VIP neurons contributing to reversal learning predominantly through RPE processing, while PV neurons do so through both reward and punishment processing. Despite the need for further clarification, it is worth noting that manipulating VIP, but not PV neuronal activity significantly prolonged the duration of reversal learning at the behavioral level. Note that direct comparisons of inactivation results are challenging due to various factors, such as the efficiency of viral gene delivery, which can influence the final outcome. Nevertheless, these results align with the idea that VIP neurons exert powerful influences on mPFC neural circuit operations through disinhibitory control^32^. As potent regulators, VIP neurons might specialize in monitoring significant changes in environmental contingencies and issuing reward-related error correction signals, thereby facilitating rapid behavioral adjustments. VIP neurons in sensory cortical areas show enhanced responsiveness to novel or unexpected sensory stimuli^35,36^, and they dynamically modulate other neurons’ sensory responses according to factors such as locomotion, behavioral strategy, and arousal^37-40^. These findings, together with our own, suggest that cortical VIP neurons may specialize in a supervisory role, wherein they oversee notable deviations from established internal and external states and intervene in ongoing cortical processes as necessary.

## Methods

### Behavioral task

Mice were trained to perform a probabilistic classical conditioning task under head fixation^21,30^ (**Fig. 1a**). All mice were habituated in the task chamber for 1-2 days before training began. We used two slightly different versions of a probabilistic classical conditioning task (tasks 1 and 2). In task 1, two out of four different odor cues (citral, 1-butanol, isoamyl acetate, and L-carvone; Sigma-Aldrich, MO, USA) were paired with a reward (2 ul of water) and the other two with a punishment (200 ms, 1.4 psi air puff delivery to the right eye) at a 75% probability. In each trial, a randomly chosen odor cue was presented for 1 s, followed by a 1 s delay before the delivery of the US (water reward or air puff). A new trial began after an inter-trial interval (ITI) ranging from 3 to 5 s, distributed uniformly (**Fig. 1a**). Mice were initially trained with a single set of CS_rw_ and CS_pn_, gradually progressing to two sets over 2-4 daily sessions. This task was employed in experiments involving chemogenetic or optogenetic manipulation.

Following successful cue discrimination (significantly higher licking in CS_rw_ than CS_pn_ trials in all CS_rw_ versus CS_pn_ comparisons in a moving analysis window of 50 trials within 300 trials since session onset; Tukey-Kramer post-hoc test following one-way ANOVA, *p* < 0.05)^21^, the reversal training phase began. Each daily session triggered reversal once the mouse reached a predefined behavioral threshold (significantly higher licking in CS_rw_ than CS_pn_ trials in all CS_rw_ versus CS_pn_ comparisons in a moving window of 50 trials; Tukey-Kramer post-hoc test following one-way ANOVA, *p* < 0.05), and after completing at least 100 trials. A randomly chosen pair of CS_rw_ and CS_pn_ had their cue-outcome contingencies switched following the onset of reversal.

In chemogenic experiments, mice underwent one episode of reversal training until they met the performance criterion (significantly higher licking in CS_rw_ than CS_pn_ trials in all CS_rw_ versus CS_pn_ comparisons (moving window of 50 trials) in fewer than 300 trials during a daily session of 600 trials; Tukey-Kramer post-hoc test following one-way ANOVA, *p* < 0.05). If the animal failed to reach the reversal criterion within a daily session, the last trial number (600) was considered as the success trial (n = 1 animal). In optogenetic experiments, mice underwent additional training to experience two episodes of reversal within a daily session. During the second reversal, a cue previously subjected to a contingency reversal in the first episode was randomly selected. Depending on its expected outcome, the cue predicting the opposite outcome and not previously subjected to a contingency reversal was selected for reversal. The second reversal occurred after 100 trials following the successful completion of the first reversal. Before optogenetic activation commenced, the mice underwent training until they successfully completed both episodes of reversal within 200 trials each. Daily sessions comprised 600 to 790 trials.

In task 2, employed for calcium imaging, one out of three different odor cues (alpha-terpinene, black cherry and citral; Sigma-Aldrich) was paired with a reward (2 ul of water), another with a punishment (air puff delivery to the left eye) at a 75% probability, and the remaining one with no outcome (CS_rw_, CS_pn_, and CSnt, respectively). Each trial involved the delivery of a randomly selected odor cue (CS) for 1.5 s, followed by a 1.5-s delay (delay 1) before advancing a water port toward the animal for licking. After an additional 1.5-s delay (delay 2), a drop of water or an air puff (US) was given with a 75% probability. The water port was retracted 2.5 s after US delivery (or omission), followed by a variable ITI (uniform random distribution; 3.5–5.5 s; **Fig. 3a**). Mice were trained in the task until their delay-2 anticipatory lick rate differed significantly across CS_rw_, CSnt, and CS_pn_ trials (significantly higher licking in CS_rw_ than CSnt and CS_pn_ trials in a moving analysis window of 40 trials within 75 trials since reversal onset; Tukey-Kramer post-hoc test following one-way ANOVA, *p* < 0.05; total 1-3 daily sessions) within a total of 420 trials per session.

Mice were then subjected to reversal learning, comprising three phases (pre-reversal, reversal, and post-reversal). Initially, the mice encountered the same CS-US contingencies as the final phase of the previous session (pre-reversal phase). Reversal occurred after a minimum of 140 trials since training onset and 75 trials since the mouse crossed the behavioral threshold (significantly higher delay-2 anticipatory licking in CS_rw_ than CSnt and CS_pn_ trials in a moving window of 40 trials; Tukey-Kramer post hoc test following one-way ANOVA, *p* < 0.05). After reversal onset, CS_rw_ was paired with 75% punishment (CS_rw→pn_), while CS_pn_ was with 75% reward (CS_pn→rw_). The animal was trained until the delay-2 lick rate met the reversal criterion (significantly higher anticipatory licking in CS_pn→rw_ than CSnt and CS_rw→pn_trials in a moving window of 40 trials; Tukey-Kramer post hoc test following one-way ANOVA, *p* < 0.05; reversal phase). Subsequently, the same contingency was maintained until trial 420 after reversal (post-reversal phase). The mice were trained until they reached the reversal criterion within 150 trials since reversal onset (2-7 daily sessions). For the analysis of VIP neuronal activity, among the final three sessions meeting the aforementioned criterion, we chose one session from each animal considering the quality of recording and behavioral performance.

### Animals

VIP-Cre knockin mice (JAX031628, Jackson Laboratory, ME, USA; n = 45, comprising 35 male and 10 female) underwent extensive handling and were water-deprived throughout the experiments, with body weights maintained at > 80% of ad libitum levels. A subset of mice (seven male and six female for Gq; three male and four female for Gi) were used for chemogenetic modulation, while others (five male for VIP neuronal perturbation and five male for its control) were allocated for optogenetic modulation, and some (12 male) for calcium imaging. They were 8-10 weeks old at the time of virus injection. Each mouse was housed individually, and all experiments were carried out during the dark phase of a 12 hr light/dark cycle. All animal care and experimental procedures were approved by the Animal Care and Use Committee of the Korea Advanced Institute of Science and Technology (KAIST) under the approval number KA2022-031.

### Surgery

The mouse was anesthetized with isoflurane (1.5 to 2.0% [vol/vol] in air) and its head was fixed in a stereotaxic frame (Kopf 940, Kopf, CA, USA). One (for calcium imaging) or two (for bilateral chemogenetic or optogenetic modulation) small burr holes (diameter, 1.5 mm for calcium imaging and 0.5 mm for chemogenetic/optogenetic modulation) were drilled into the skull (mPFC: 1.80 mm anterior and 0.33 mm lateral), targeting the mPFC. Target virus was injected into the mPFC, approximately 1.5 mm below the brain surface, at a rate of 0.05 μl/min. For calcium imaging, following virus injection, a GRIN lens with a prism (diameter, 1 mm, length, 4.3 mm, 1 mm × 1 mm prism attached, 1050-004601, Inscopix, CA, USA) was implanted, targeting layers 2/3 in the mPFC (2.0 mm ventral to brain surface). If necessary, the scalp skin at the injection site was sutured or cemented to aid in postoperative recovery. At least one week was allowed for recovery from surgery before commencing experiments.

### Chemogenetics

For chemogenetic activation or inactivation of VIP neurons, Cre-dependent AAV carrying hM3D(Gq) or hM4D(Gi) (pAAV2-hSyn-DIO-hM3D(Gq)-mCherry, Addgene plasmid #44361, MA, USA; pAAV2-hSyn-DIO-hM4D(Gi)-mCherry, Addgene plasmid #44362) was injected bilaterally in the mPFC. CNO (C0832, Sigma-Aldrich) was initially dissolved in DMSO (10 mg/ml, D2650, Sigma-Aldrich) and then diluted in PBS (0.5 ml) and administered by intraperitoneal injection (0.5 mg/kg in PBS) 30-40 min prior to the behavior session on the test day.

### Optogenetics

The AAV carrying the gene for ChRmine (AAV9-Ef1a-DIO-ChRmine-mScarlet-Kv2.1-WPRE, Addgen plasmid #130999) or tdTomato (AAV2-CAG-FLEX-tdTomato, UNC Vector Core, NC, USA; control) was injected bilaterally in the mPFC. Transcranial opto-stimulation involved the attachment of an optic fiber on the mouse’s skull between the two virus injection sites. Light stimulation (635 nm; frequency, 20 Hz; duration, 5 ms; intensity, 800 mW mm^-2^) was delivered using a PhoxX laser diode module (Omicron, Rodgau, Germany). Stimulation occurred during the outcome period, spanning from 0.5 s before to 2 s after outcome onset. The stimulation initiated at the onset of either the first or the second reversal episode, alternating across daily sessions, and concluded when the animal met the reversal criterion. To mitigate the mice’s perception of the light flashes, the experiments were conducted under red light.

### Calcium imaging

The AAV carrying the gene for a genetically encoded calcium indicator (pGP-AAV1-syn-FLEX-jGCaMP7f-WPRE, Addgene plasmid #104492) was injected unilaterally into the right mPFC (1.5 mm ventral from brain surface) of VIP-Cre mice. A gradient refractive index (GRIN) lens with an attached prism was implanted approximately 0.1 mm lateral to layer 2/3 of the prelimbic cortex. The implantation site was located at 1.80 mm anterior and 0.33 mm lateral to bregma, with a depth of 1.8 mm ventral to the brain surface. This placement targeted VIP neurons within the region spanning 1.34-2.31 mm anterior to bregma and 1-2 mm ventral to the brain surface.

Following a 6-8 week period since lens implantation, the baseplate for the nVista microscope was secured to the skull. Prior to baseplate placement, calcium signals were evaluated to optimize the positioning of the nVista 3.0 microscope (Inscopix). Calcium signals were recorded using the implanted GRIN lens and nVista microscope, capturing data at a frame rate of 30 Hz with an LED power of 0.4-0.6 mW/mm^2^. The acquired calcium imaging data underwent spatial downsampling (1/4 scale) and motion correction using Inscopix Data Processing Software (version 1.3.1; Inscopix) and turboreg^41^. The processed video was exported in TIFF format and further analyzed using the CNMF-E algorithm^42^ to extract single unit signals. Only units with mean event rates ≥ 0.025 Hz were included in the analysis.

### Histology

Following the conclusion of calcium imaging or behavioral experiment, the mouse brain was perfused with 10 ml of cold 0.1 M PBS, followed by the addition of 10 ml of ice-cold 4% PFA using a 10-ml syringe. The mouse brain was initially incubated in 4% PFA at 4°C for 7 days, after which the lens was gently removed from the brain. The brain was then incubated in 30% sucrose in 0.1 M PBS at 4°C for 2 days. The fixed, dehydrated brain was cut into 40-μm-thick coronal slices using a cryostat (Leica CM3050 S; Leica Biosystems, Wetzlar, Germany) and washed with 0.1 M PBS for 10 minutes. These slices were mounted with 4’,6-diamidino-2-phenylindole (DAPI)-containing Vectashield (Vector Laboratories, CA, USA) and then imaged on a slide-scanning microscope with a 10×/0.45NA objective using Zen 2.5 software (blue edition, Axio scan Z1; Zeiss, Jena, Germany) or a confocal microscope with 10×/0.45NA, 20×/0.8NA, and 40×/1.20NA water objectives using Zen 2011 SP7 software (black edition, LSM 780; Zeiss). The images were processed using Zen 2.5 lite software (blue edition; Zeiss).

### Analysis

#### Response normalization

Unless noted otherwise, calcium transient signals were normalized by calculating the z-score relative to the mean and SD of the baseline period (0.5-s period before cue onset) activity across trials (baseline normalization). However, for the analyses related to the detection of reversal onset, calcium transient signals during a given analysis time window were normalized across trials within 100 trials before reversal onset (trial normalization; **Fig. 5, Supplementary Fig. 3**, and **Supplementary Fig. 4**).

#### Cue and outcome-preference indices

Cue-preference index was defined as (*A*-*B*), where *A* and *B* are mean baseline-normalized calcium activity during the given analysis time window in CS_rw_ and CS_pn_ trials (for neural activity before the reversal) or CS_rw→pn_ and CS_pn→rw_ (for neural activity after reversal) trials, respectively. Reward-preference index was defined as (*C*-*D*), where *C* and *D* are mean baseline-normalized calcium activity during the 1.5-s time window since the first lick following delay offset in reward trials and since delay 2 offset in reward-omission trials, respectively. Punishment-preference index was defined as (*E*-*F*), where *E* and *F* are mean baseline-normalized calcium activity during the 1.5-s time window since delay 2 offset in punishment and punishment-omission trials, respectively.

#### Responses to task events

Neuronal responses to odor cue, lick-port advancement, and ITI onsets were assessed by calculating an event modulation index for each of these events as (*After*-*Before*), where *Before* and *After* denote mean baseline-normalized calcium transients during 0.5-s time window before and after each event onset.

#### Response to reversal onset

To examine dynamics of cue-dependent VIP neuronal activity related to monitoring reversal onset, we analyzed neural activity during the first delay period (delay-1 activity; 1.5 s) using trials around reversal onset. We chose four trials, each presenting a specific cue (CS_rw→pn_ or CS_pn→rw_), both before and after encountering the first unexpected outcome (the first outcome delivery since reversal onset). We then normalized delay-1 activity in these trials with the mean and SD across CS_rw→pn_ and CS_pn→rw_ trials within 100 trials before reversal onset (trial normalization) and the responses to CS_rw→pn_ and CS_pn→rw_ in the corresponding ordinal positions were averaged. We also analyzed PP and FS neurons from our previous study^21^ in a similar manner, except that neural activity during the 1-s delay period was analyzed. Only those units with mean firing rates ≥ 1 Hz during the delay period were included in the analysis. For outcome responses of VIP neurons, four trials both before and after the reversal onset with reward or punishment presentation were chosen. The outcome period activity (1.5 s) was normalized with the mean and SD across outcome-delivery trials within 100 trials before reversal onset.

#### Multiple regression analysis

Neural signals for reward (*Rw(t)*) during the outcome period (first 1.5 s window) of VIP neurons were analyzed using only CS_rw_ trials with the following multiple linear regression model:

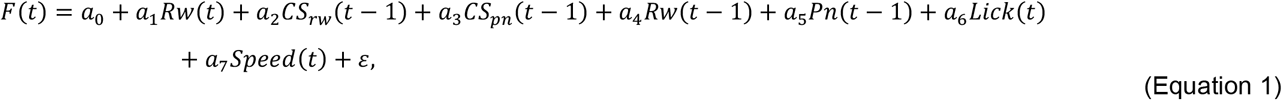

Punishment signals (*Pn(t)*) during the outcome period were analyzed using only CS_pn_ trials with the following multiple linear regression model:

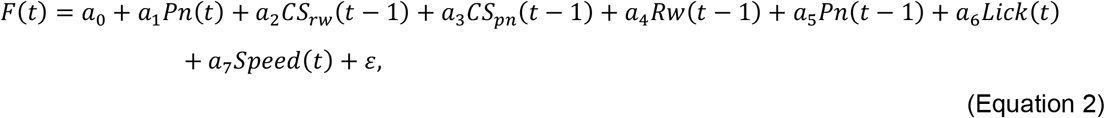

Neural signals for CS (*CS*_*rw*_*(t)* and *CS*_*pn*_*(t)*) during the outcome period were analyzed using all trials with the following multiple linear regression model:

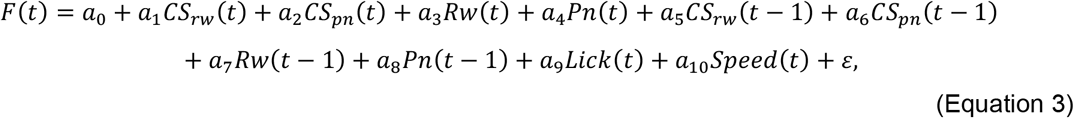

where *F(t)* represents a calcium transient trace during the outcome period, *a0* to *a10* are regression coefficients, *CS*_*rw*_*(t), CS*_*pn*_*(t), Rw(t)*, and *Pn(t)* represent reward predicting cue, punishment predicting cue, reward, and punishment (1 if present and 0 otherwise), respectively, *Lick(t)* and *Speed(t)* are lick numbers and running speed, respectively, during outcome period in trial *t*, and *ε* is the error term.

For PP and FS neurons, the following multiple linear regression model was used to analyze reward (*Rw(t)*) neural signals during the outcome period (first 1.5 s window) using only CS_rw_ trials:

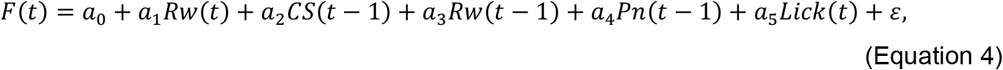

Punishme nt signals (*Pn(t)*) were analyzed with the following multiple linear regression model using only CS_pn_ trials:

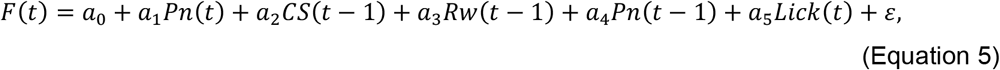

For CS signals (*CS(t)*) during the outcome period, the following multiple linear regression model was used:

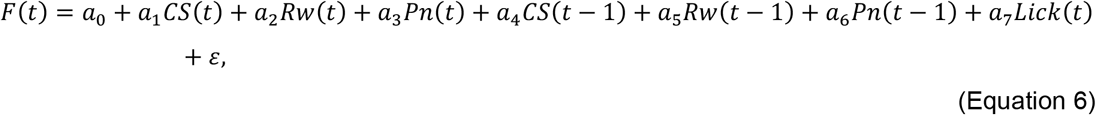

where *F(t)* represents neural firing rate during the outcome period, *a0* to *a7* are regression coefficients, *CS(t), Rw(t)*, and *Pn(t)* represent trial cue (1 for CS_rw_ and -1 for CS_pn_), reward and punishment (1 if present and 0 otherwise), respectively, *Lick(t)* is lick numbers during outcome period in trial *t*, and *ε* is the error term. Only those units with mean firing rates ≥ 1 Hz during the outcome period were included in the analysis.

#### Analysis of RPE signals

To examine neural activity related to RPE, we calculated the Pearson’s correlation coefficient between population *t*-values of *CS*_*rw*_*(t)* and *Rw(t)* terms (*r*_*CS-Rw*_) of the multiple regression model (eq 2, 3; **Fig. 6a**). A significant (*p* < 0.05) negative correlation was considered indicative of RPE signals for a given neural population^30^. To assess the dynamics of RPE-related neural activity during reversal learning, we constructed the temporal profile of *r*_*CS-Rw*_ over trials with a 100-trial sliding window (**Fig. 6b-c**). This analysis was conducted for all analyzed neurons to evaluate a general trend of RPE signal dynamics (100-trial moving analysis window, 10-trial steps; **Fig. 6b**), as well as for simultaneously recorded neurons from each animal (100-trial moving analysis window, 1-trial steps; **Fig. 6c**).

To determine the persistence of RPE signals for each animal, we first smoothed the *r*_*CS-Rw*_ trace by applying a 40-trial moving average to mitigate the impact of noisy fluctuations. We then calculated RPE persistence by determining the step difference between the trial where *r*_*CS-Rw*_ is minimal within 93 trials (the mean number of trials to reach the reversal criterion) since reversal onset and the trial where *r*_*CS-Rw*_ is maximal since reversal onset. Only mice with more than three units with mean calcium event rates ≥ 0.025 Hz were included in this analysis (n = 10; two mice excluded from the analysis).

To compare the time courses of RPE signals and behavioral performance, we constructed the temporal profile of LDI. LDI was defined as (*G*-*H*)/(*G*+*H*), where *G* and *H* are mean anticipatory lick numbers in CS_rw→pn_ and CS_pn→rw_ trials, respectively, during the first delay period, for each animal using a 100-trial moving analysis window in 1-trial steps. We then determined saturation points for both LDI and *r*_*CS-Rw*_ traces. The saturation point is the trial where LDI (or *r*_*CS-Rw*_) crosses 5% (or 95%) value between the maximum and minimum values (**Fig. 6c**).

Temporal dynamics of *r*_*CS-Rw*_ was further examined by fitting the *r*_*CS-Rw*_ trace to the following sigmoidal function^21^ (**Supplementary Fig. 7a-c**) using ‘fit’ function of MATLAB (MathWorks Inc., MA, USA):

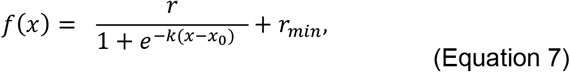

where *x0* is the midpoint trial, *k* is the steepness, *r* is the range, and *rmin* is the minimum value. Only animals with *R*^*2*^ > 0.4 were included in the analysis (n = 8; four mice excluded from the analysis). The saturation point of the fitted sigmoidal function was calculated using the following equation^45^:

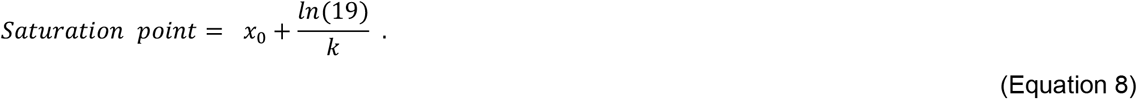

#### ANCCR model

To complement RPE-related analysis, we also analyzed VIP neuronal data using a recently proposed retrospective learning model (ANCCR^31^). The analysis was conducted using the codes provided by the authors on the repository site (https://github.com/namboodirilab/ANCCR). Three parameters, α (learning rate), R_pn_ (punishment magnitude), and sampling period, were determined for each animal (n = 12; all mice were included in the analysis) to maximize the correlation between anticipatory lick numbers and the sum of multiplications of causal weights and successor representation contingency (representing the prospective association strength). Other parameters were set as in Jeong et al. (2022)^31^. In order to accommodate rapid cue discrimination observed during the pre-reversal phase, which is attributable to the prior day’s experience, we analyzed behavioral data extending from the reversal onset on the preceding day’s session to the conclusion of the reversal session for parameter selection. The parameter selection was done using the ‘fminsearch’ function in MATLAB, whereby the best parameter set was obtained from 100 independent searches conducted with random initial parameters between 0 and 1^46^.

Neural signals for the retrospective association between CS_rw_ and reward during the outcome period (1.5 s), calculated with the ANCCR model, were examined with the following multiple linear regression model:

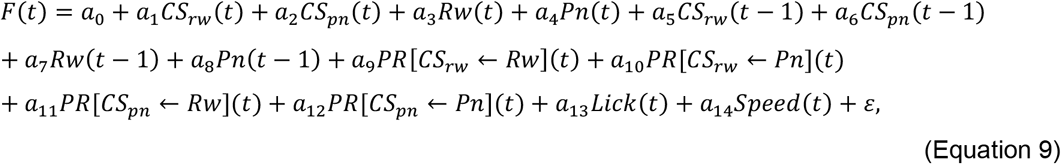

where *PR[CS*_*rw*_*←Rw](t)* represents the predecessor representation (PR) value of CS_rw_ over reward (Rw) in trial *t*.

To determine PR[CS_rw_←Rw] signaling dynamics during reversal, we constructed the temporal profile of *PR[CS*_*rw*_*←Rw](t)* absolute *t*-value using a 100-trial analysis window (1-trial step) over trials for each animal. We smoothed the trace by applying a 40-trial moving average to mitigate the impact of noisy fluctuations. We then determined PR[CS_rw_←Rw] persistence by determining the step difference between the trial where *PR[CS*_*rw*_*←Rw](t)* absolute *t*-value is maximal within 93 trials (the mean number of trials to reach the reversal criterion) since reversal onset and the trial where it is minimal since reversal onset. The saturation point was determined as the trial where *PR[CS*_*rw*_*←Rw](t)* absolute *t*-value crosses 5% value between the maximum and minimum values since reversal onset (**Fig. 6f**).

#### Statistical analysis

All statistical tests were performed with MATLAB (version R2017a). Student’s *t*-test, one-way ANOVA, Turkey-Kramer post-hoc test, one-way repeated measures ANOVA, Bonferroni’s post-hoc test, and two-way repeated measures ANOVA were used for group comparison. All tests were two-tailed, and *p*-value < 0.05 was used as the criterion for a significant statistical difference. Results are presented as mean±SEM unless noted otherwise.

## Notes

### Competing Interest Statement

The authors have declared no competing interest.

