## Supplementary Figs. 1-9, Supplementary Table 1 for "Selective engagement of prefrontal VIP neurons in reversal learning"

This PDF file includes:

Supplementary Figures 1 to 9

and Supplementary Table 1.

*These authors contributed equally to this work.

**Supplementary Figures**

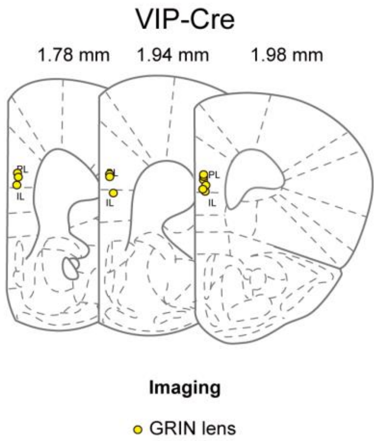


**Supplementary Fig. 1 | Histological localization of GRIN lens positions.** Shown are coronal section views of the mouse brain (left to right, 1.78, 1.94, and 1.98 mm anterior to bregma). Yellow circles indicate locations of prism tips affixed to GRIN lenses used for calcium imaging of VIP neurons (12 VIP-Cre mice). Related to Fig. 3.

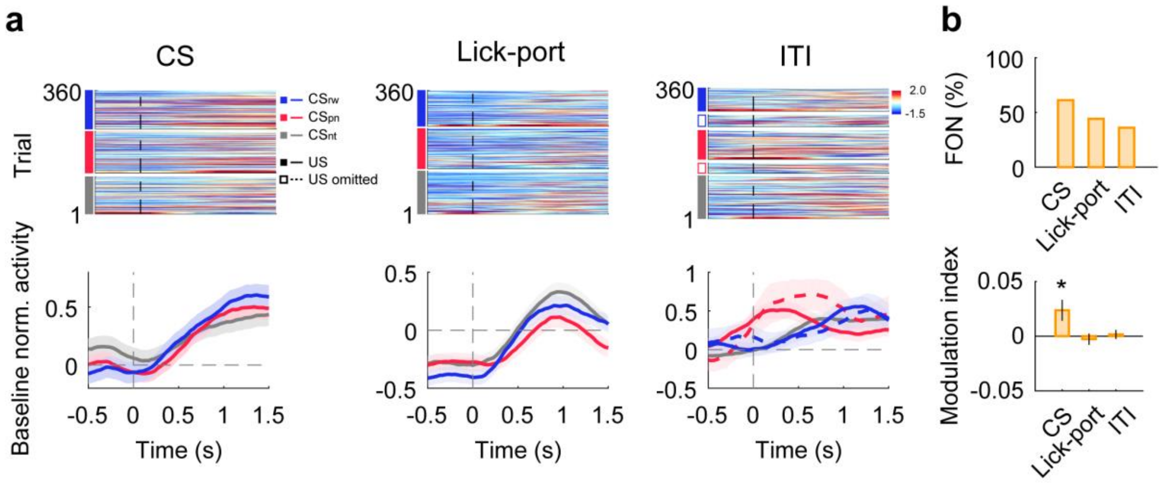


**Supplementary Fig. 2 | VIP neurons are responsive to diverse task events. (a)** Sample neurons showing differential firing between before and after the onset of CS, lick-port advancement, or ITI. The same format as in Fig. 4b. **(b)** Upper, fractions of neurons (FON) significantly responsive to CS, lick-port advancement, or ITI onset ($p < 0.05$, $t$-test comparing 0.5-s windows before and after event onset). Lower, mean (±SEM across 111 neurons) event modulation index (positive and negative, higher and lower firing after than before event onset, respectively; see Methods). *$p < 0.05$, difference from zero, $t$-test. Related to Fig. 3.

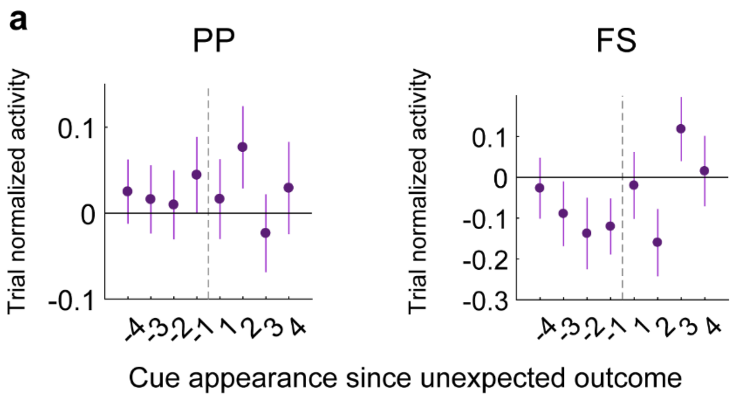


**Supplementary Fig. 3 | Cue responses of pyramidal and fast spiking neurons around reversal onset.** Shown are trial-normalized responses of putative pyramidal (PP, **a**) and fast spiking (FS, **b**) neurons during the delay period (1.0 s; FS, n = 67; PP, n = 310 neurons recorded from six animals). The same format as in Fig. 5b. Related to Fig. 5.

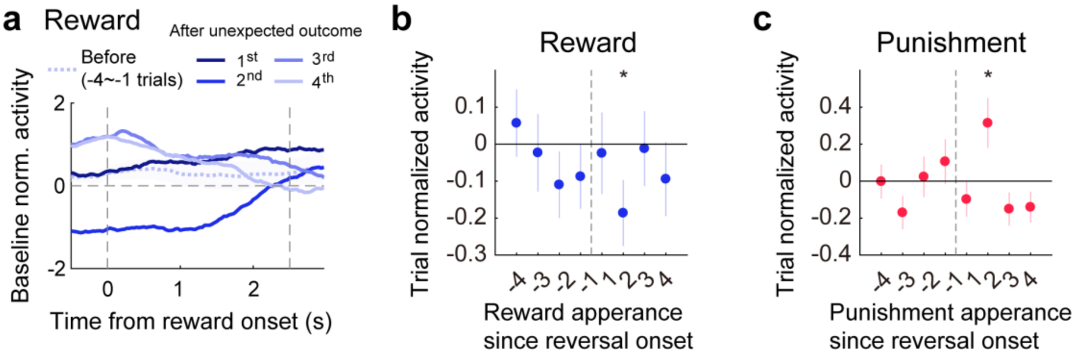


**Supplementary Fig. 4 | Outcome-period responses of VIP neurons around reversal onset. (a)** Baseline-normalized outcome responses of a sample VIP neuron to reward delivery before (averaged over 4 trials) and after reversal onset. Note diminished neural activity preceding outcome onset in the second trial, which can be attributed to decreased response to the first encounter of $CS_{rw}$ since unexpected outcome delivery (Fig. 5). **(b)** Group data showing trial-normalized outcome-period responses to reward delivery around reversal onset. **(c)** Similar to (b), responses to punishment delivery are depicted. *$p < 0.05$, difference from zero, *t*-test (n = 106 neurons recorded from 12 animals). The same format as in Fig. 5 except that outcome-period responses were analyzed instead of the first delay-period responses. Related to Fig. 5.

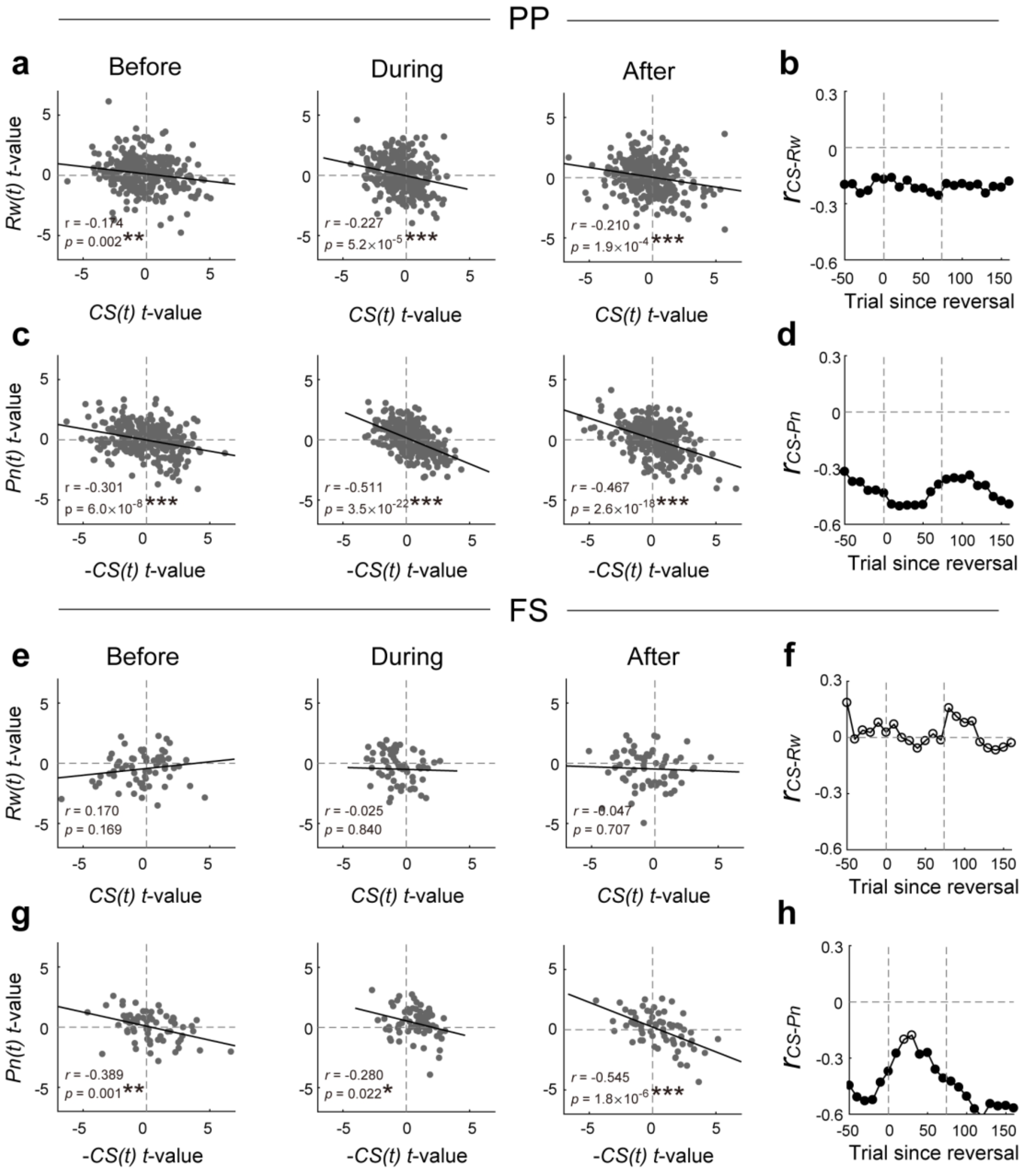


**Supplementary Fig. 5 | Prediction error-related activity of pyramidal and fast spiking neurons.** We analyzed PP and FS neuronal data from our previous study[21] using multiple regression (eq. 4-6). The scatter plots depict CS-related (abscissa) and outcome-related (ordinate) neural activity during the outcome-period (1.5 s). Since the study involved only two odor cues ($CS_{rw\rightarrow pn}$ and $CS_{pn\rightarrow rw}$), CS-related neuronal activity was represented by a single regression term (*CS(t)*; 1 and -1 denoting reward- and punishment-predicting cue, respectively) in the multiple regression model (eq. 6). Therefore, positive and negative *t*-values for *CS(t)* correspond to *t*-values for $CS_{rw}(t)$ and $CS_{pn}(t)$, respectively, in VIP neural data. **(a to d)** Results from the analysis of PP neuronal activity (n = 306 recorded from six animals) regarding reward prediction-related activity **(a and b)** and punishment prediction-related activity **(c and d).** The same format as in Fig. 6a-b. **(e to h)** Results from FS neurons (n = 67 recorded from six animals). The same format as in a-d. *$p < 0.05$, **$p < 0.01$, ***$p < 0.001$. Related to Fig. 6.

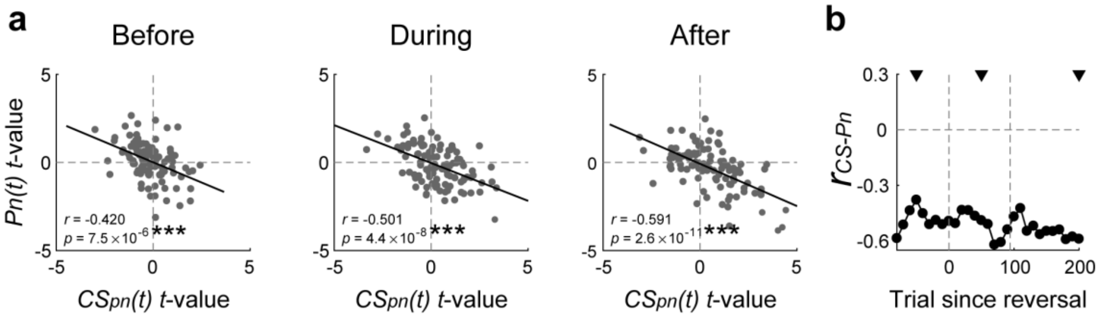


**Supplementary Fig. 6 | VIP neurons encode punishment prediction error across all stages of reversal learning.** Outcome-period VIP neuronal responses to punishment-predicting cue ($CS_{pn}(t)$; $\mathrm{CS}_{\mathrm{pn}\rightarrow\mathrm{rw}}$ and $\mathrm{CS}_{\mathrm{rw}\rightarrow\mathrm{pn}}$ before and after reversal, respectively) and punishment (*Pn(t)*) were analyzed using multiple regression as in Fig. 6. **(a and b)** The same format as in Fig. 6a-b. ***$p < 0.001$. Related to Fig. 6.

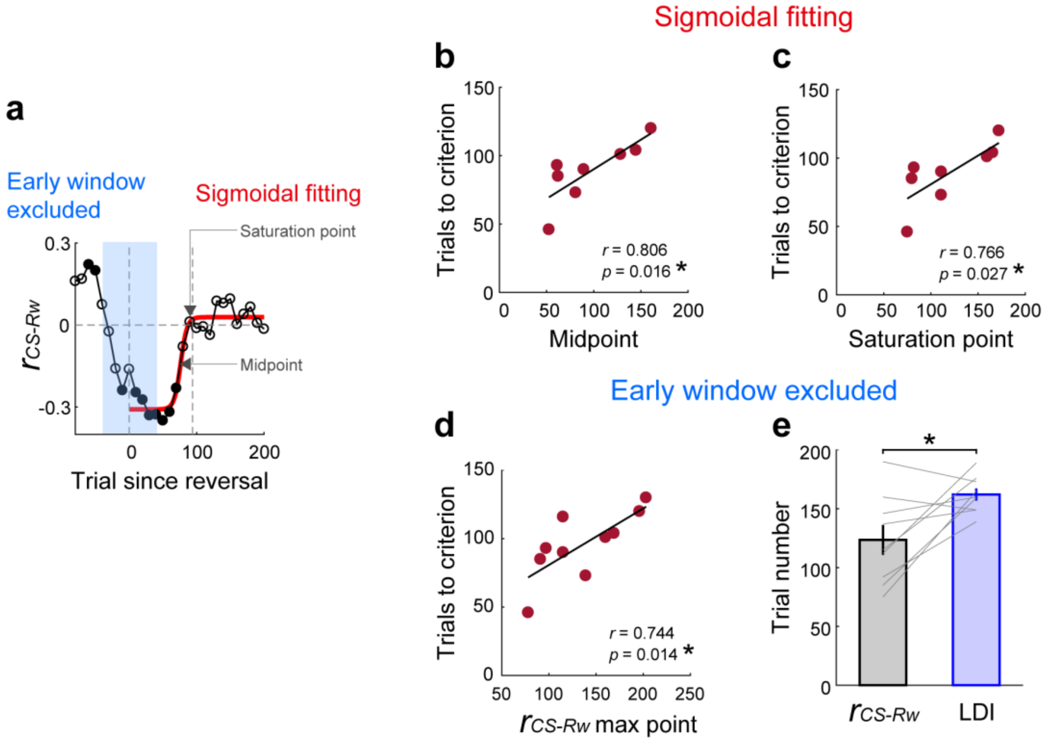


**Supplementary Fig. 7 | Additional analysis results related to VIP neuronal error correction signals. (a)** A schematic showing sigmoidal fitting (red line) of $r_{CS\text{-}Rw}$ dynamics after reversal onset (the same graph shown in Fig. 6b). Arrows indicate the midpoint and saturation point of the sigmoidal curve. **(b and c)** Similar to the analysis shown in Fig. 6d, with the midpoint (b) or saturation point (c) of the sigmoidal curve used as an index for the persistence of RPE signals (n = 8 mice with $R^2 > 0.4$ for the sigmoidal fitting). **(d)** Early analysis windows (0-50 trials since reversal onset) including pre-reversal trials were excluded from the analysis to prevent pre-reversal period activity from affecting RPE persistence estimation. The abscissa ($r_{CS\text{-}Rw}$ max point) denotes the trial where the value of $r_{CS\text{-}Rw}$ is maximal. The same format as in Fig. 6d. **(e)** Trial numbers for $r_{CS\text{-}Rw}$ max point (black) and LDI saturation point (blue), excluding the early analysis windows. Gray lines, individual animal data (n = 10 mice). Bar graphs, their mean and SEM. *$p < 0.05$, *t*-test. Related to Fig. 6.

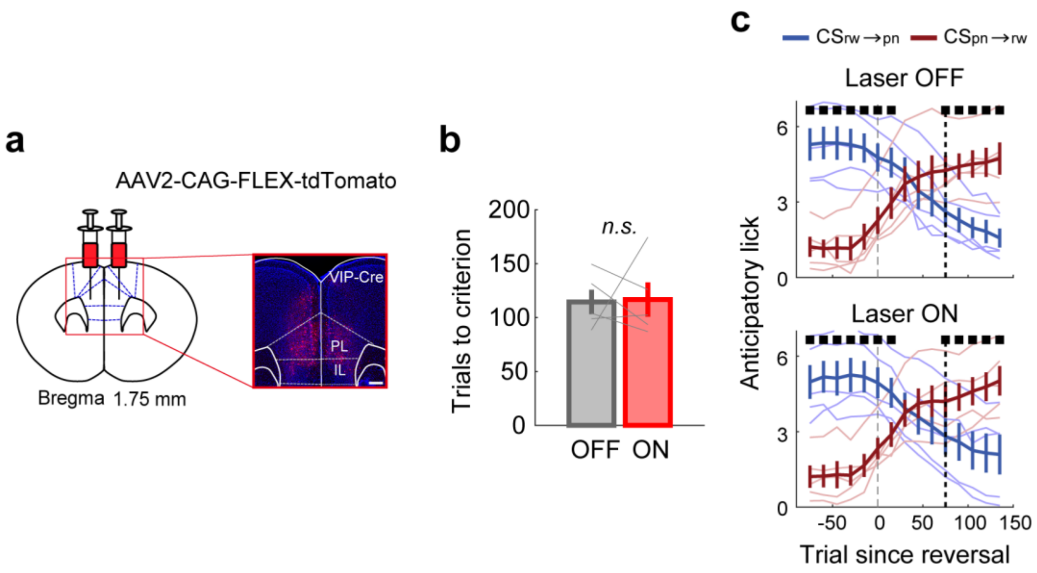


**Supplementary Fig. 8 | Optogenetic modulation has no significant effect on reversal learning in control animals. (a to c)** The same format as in Fig. 7. Error bars, SEM across animals (n = 5). Related to Fig. 7.

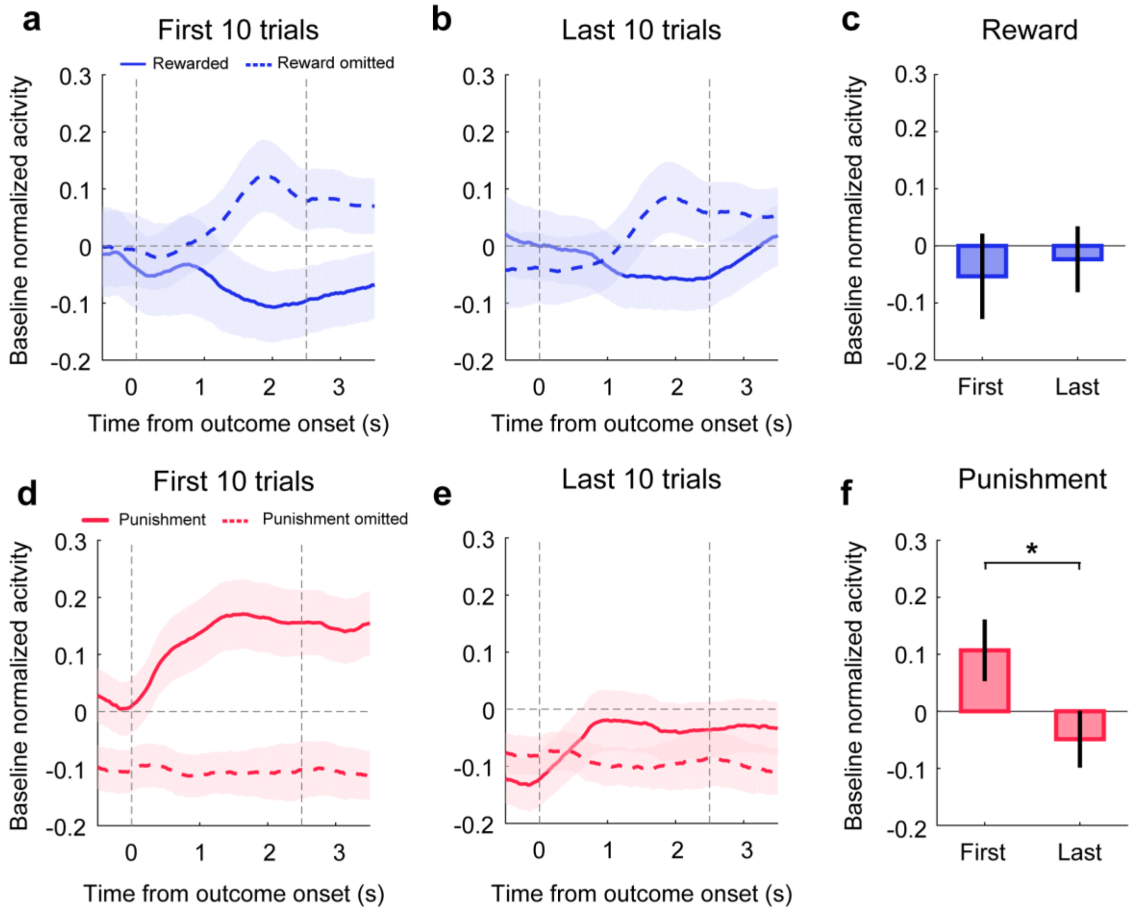


**Supplementary Fig. 9 | Outcome responses of VIP neurons during early and late trials**. **(a to b)** Mean baseline-normalized VIP neuronal responses to reward delivery (solid line) and omission (dashed line) during the first (a) and last (b) 10 trials during the pre-reversal phase of the reversal session (n = 106 VIP neurons). **(c)** Mean normalized responses to reward delivery during 1.5 s after reward onset. **(d to f)** Results for the punishment responses of VIP neurons. The same format as in a to c. Shading and error bars, SEM across 106 VIP neurons. *$p < 0.05$, $t$-test.

**Supplementary Table 1 | The numbers of mice and neurons analyzed in each experiment.**

| Figures | Experiment | Mouse genotype and number | Number of neurons |
|---|---|---|---|
| Fig. 1, 2 | Chemogenetic modulation of VIP neurons | VIP-Cre (Gq, 7 male and 6 female; Gi, 3 male and 4 female) | |
| Fig. 3<br>Ex. Fig. 2 | VIP neuronal calcium imaging | VIP-Cre (Before reversal training, 9 male) | n = 111 |
| Fig. 4,5,6<br>Ex. Fig. 4,5,7,9 | VIP neuronal calcium imaging | VIP-Cre (Reversal training, 12 male) | n = 106 |
| Fig. 7<br>Ext.Fig. 8 | Optogenetic modulation of VIP neurons | VIP-Cre (ChRmine, 5 male; tdTomato, 5 male) | |
| Ext.Fig. 3 | Single unit recording with tetrodes | PV-Cre (2 male) & SST-Cre (4 male) | PP, n = 310; FS, n = 67 |
| Ext.Fig. 6 | Single unit recording with tetrodes | PV-Cre (2 male) & SST-Cre (4 male) | PP, n = 306; FS, n = 67 |
